# VRK1 is required in VRK2-methylated cancers of the nervous system

**DOI:** 10.1101/2021.12.28.474386

**Authors:** Jonathan So, Nathaniel W. Mabe, Bernhard Englinger, Sydney M. Moyer, Maria C. Trissal, Joana G. Marques, Jason Kwon, Brian Shim, Eshini Panditharatna, Daeun Jeong, David Mayhew, Justin Hwang, Kimberly Stegmaier, Mariella G. Filbin, William C. Hahn

## Abstract

Collateral lethality occurs when loss of one paralog renders cancer cells dependent on the remaining paralog. Combining genome scale CRISPR/Cas9 screens coupled with RNA-sequencing in over 900 cancer cell lines, we found that cancers of nervous system lineage, including adult and pediatric gliomas and neuroblastomas, required the nuclear kinase Vaccinia-Related Kinase 1 (VRK1) for their survival. *VRK1* dependency was inversely correlated with expression of its paralog VRK2. *VRK2* knockout (KO) sensitized cells to *VRK1* suppression, and conversely, VRK2 overexpression increased cell fitness in the setting of VRK1 suppression. DNA methylation of the *VRK2* promoter was associated with low VRK2 expression in human neuroblastomas, and adult and pediatric gliomas. Mechanistically, depletion of *VRK1* reduced Barrier-to-Autointegration Factor (BAF) phosphorylation during mitosis, resulting in DNA damage and apoptosis. Together, these studies identify VRK1 as a synthetic lethal target in *VRK2* promoter-methylated adult and pediatric gliomas and neuroblastomas.

**Statement of Significance:** We credential VRK1 as a target in adult and pediatric gliomas, and neuroblastomas with *VRK2* promoter methylation. This demonstrates the utility of paralog-driven synthetic lethal interactions for biomarker-linked, targeted therapeutics.

## Introduction

Tumors of the nervous system constitute some of the most devastating malignancies in both adult and pediatric patients (1). Tumors arising in both the central and peripheral nervous system (CNS and PNS, respectively) often exhibit an aggressive clinical course and are refractory to currently available systemic therapy (2,3).

Glioblastoma is the most common primary brain tumor in adults and is characterized by poor prognosis and low cure rates (4). Diffuse midline gliomas (DMG) that harbor histone 3 lysine27-to-methionine mutations (“pediatric diffuse midline glioma, H3 K27-altered”) occur in children with a peak incidence of 6 to 9 years of age (5). Due to their infiltrative growth pattern, these gliomas are unresectable and uniformly fatal. Neuroblastoma (NB) is the most common extracranial solid tumor malignancy in childhood, commonly originating in the adrenal medulla or paraspinal ganglia (2). Standard of care treatment schemes for these tumors remains cytotoxic radio-chemotherapy and surgery, and patient prognosis has not substantially improved over the last decade (3,6–7). Thus, there is an urgent need for the identification of novel, targetable biomarkers in these tumor entities to translate into improved patient outcomes.

The implementation of CRISPR/Cas9 screening technologies has facilitated systematic studies to identify novel therapeutic targets and biomarkers of response across many cancers. Such dependency maps have unveiled a number of gene targets beyond known oncogenic drivers and hold the potential for tumor-specific, personalized therapy (8). Integration of genome-wide functional studies with genome-wide transcriptomics or epigenomics allows for correlative connections between gene dependencies and cancer transcriptional landscapes.

By comprehensively integrating genome scale, loss-of-function genetic screens, RNA-sequencing and analysis of DNA methylation patterns, we identified the nuclear, serine-threonine kinase, Vaccinia-Related Kinase 1 (VRK1), as a highly selective dependency in adult and pediatric CNS and PNS tumors that exhibit low expression of the *VRK1* paralog *VRK2*.

## Results

### VRK1 is a selective dependency in adult and pediatric glioma and neuroblastoma

The Cancer Dependency Map includes CRISPR-Cas9 loss-of-function screens performed in over 900 cell lines and 25 different cancer lineages (9). Using this dataset, we found that *VRK1* is a strong genetic dependency in adult glioma (*p* = 2×10^−12^; Student’s *t*-Test; *n* = 61) and pediatric neuroblastoma (*p* = 3×10^−8^; Student’s *t*-Test; *n* = 20) cell lines **(Fig. 1A)**. Indeed, *VRK1* was the gene with the most differential dependency in central nervous system (CNS) and peripheral nervous system (PNS) lineages **(Fig. 1B)**.

**Figure 1.**
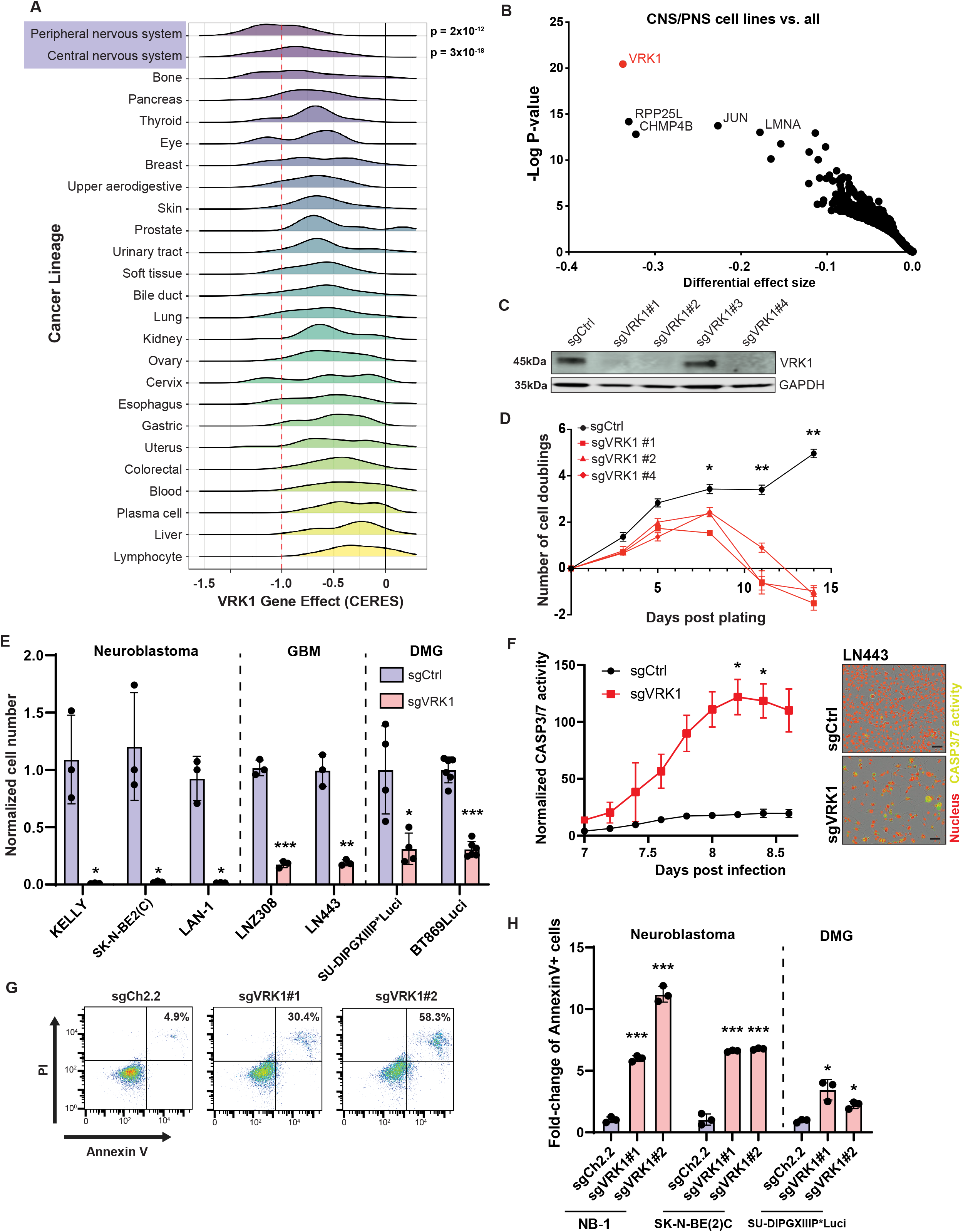
VRK1 is a dependency in GBM, NB, and DMG. **A.** Histogram plots showing *VRK1* CERES dependency scores in over 900 cell lines, representing 27 different cancer lineages from the DepMap dataset (21Q3). Compared to all other lineages cell lines in the CNS (p = 2×10^−12^) and PNS (p = 3×10^−18^), lineages were significantly more dependent on *VRK1*. **B.** Differential dependency of gene KO in CNS and PNS cell lines versus all other lineages. Gene effect size is calculated as the difference in average CERES score between lineage groupings and q-value is determined by two-class comparison with the Limma eBayes methodology. The top enriched dependencies in CNS/PNS lineages are annotated. **C.** VRK1 protein expression following expression of 4 different sgRNA in the NB-1 neuroblastoma cell line. The top three guides with greatest *VRK1* loss were carried forward in subsequent experiments. **D.** Cell proliferation following *VRK1* KO with three separate guides in NB-1 cells. sgCtrl represents a non-targeting control guide (n=3; mean ± SD). Significance at each time point was determined by two-way ANOVA (treatment x time). *p < 0.05, **p < 0.001. **E.** Single guide (sg)VRK1 KO after 14 days in cell lines representing NB (n=3), GBM (n=2), and DMG (n=2) lines. (n ≥ 3; mean ± SD plotted). **F.** Left panel: Time-course of CASP3/7 activity, as measured by cleavage of a peptide reporter, following *VRK1* KO in LN443 cells (n=3; mean ± SD). Total reporter fluorescent signal is normalized by cell confluence. Right panel: Representative images from live-cell experiment 8 days following infection with sgVRK1 or sgCtrl guide in LN443 GBM cells. Red: nuclear stain; Green: CASP3/7 activity (Incucyte Caspase-3/7 dye). Significance at each time point was determined by two-way ANOVA (treatment x time). *p < 0.05. Scale bar: 20 μm. **G.** Representative flow cytometry gating strategy for Propidium Iodide and Annexin-V staining following *VRK1* KO in NB-1 NB cells. **H.** Quantification of Annexin-V positive cells following *VRK1* KO with two separate guides in three cell lines representing NB and DMG lineages after 7 days. (n=3; mean ± SD; from 2 separate experiments). *p < 0.05, **p < 0.001, ***p < 0.0001; Two-tailed, Student’s T-test for all comparisons unless otherwise specified.

We validated that VRK1 is required for cell proliferation using CRISPR/Cas9 KO (KO). Three of the four *VRK1* sgRNAs that were included in the Cancer Dependency Map led to robust *VRK1* KO and proliferation defects in NB and GBM cultures **(Fig. 1C, Supplementary Fig. S1A-C)**. *VRK1* KO resulted in a significant decrease in cell fitness approximately 8 days following viral transduction in NB-1 cells **(Fig. 1D)**, as well as in a larger panel of neuroblastoma and GBM cell models **(Fig. 1E)**. In addition, the Cancer Dependency Map includes a pediatric glioma model (KNS42), which also demonstrated strong *VRK1* dependency. We therefore tested primary, pediatric H3K27M DMG neurosphere models and confirmed that *VRK1* singlegene KO significantly decreased cell viability **(Fig. 1E)**. To determine whether the reduced viability was due to apoptosis, we performed live-cell experiments with a CASP3/7 cleavage reporter in the LN443 GBM cell line and found significantly higher CASP3/7 activity after *VRK1* KO **(Fig. 1F)**. In an orthogonal approach, we found significant induction of apoptosis in NB and DMG models following *VRK1* KO, as assessed by Annexin-V / PI staining **(Fig. 1G, H)**. In contrast, we failed to observe significantly altered cell-cycle profiles in response to *VRK1* KO in GBM or DMG models, and only a small increase of cells in G2/M phase in NB cells **(Supplementary Fig. S2A-D)**. Taken together, these observations demonstrate that *VRK1* is a robust dependency in tumors of nervous system lineages, and *VRK1* KO results in apoptotic cell death.

### VRK2 expression is a biomarker for VRK1 dependency

To identify genes or pathways that predict *VRK1* dependency, we correlated gene expression from the Cancer Cell Line Encyclopedia (CCLE) with *VRK1* dependency and found that it is most strongly correlated with the loss of expression of its paralog *VRK2* (Pearson Correlation = 0.37, q < 10-^25^) **(Fig. 2A)**. Low *VRK2* expression was associated with *VRK2* promoter CpG methylation and was highly enriched in the CNS and PNS lineages **(Fig. 2B)**. Interestingly, gene expression data from healthy neural tissue also demonstrated low VRK2 expression relative to other tissues, suggesting *VRK2* promoter methylation may be specific to the neural lineage **(Supplementary Fig. S3A)**.

**Figure 2.**
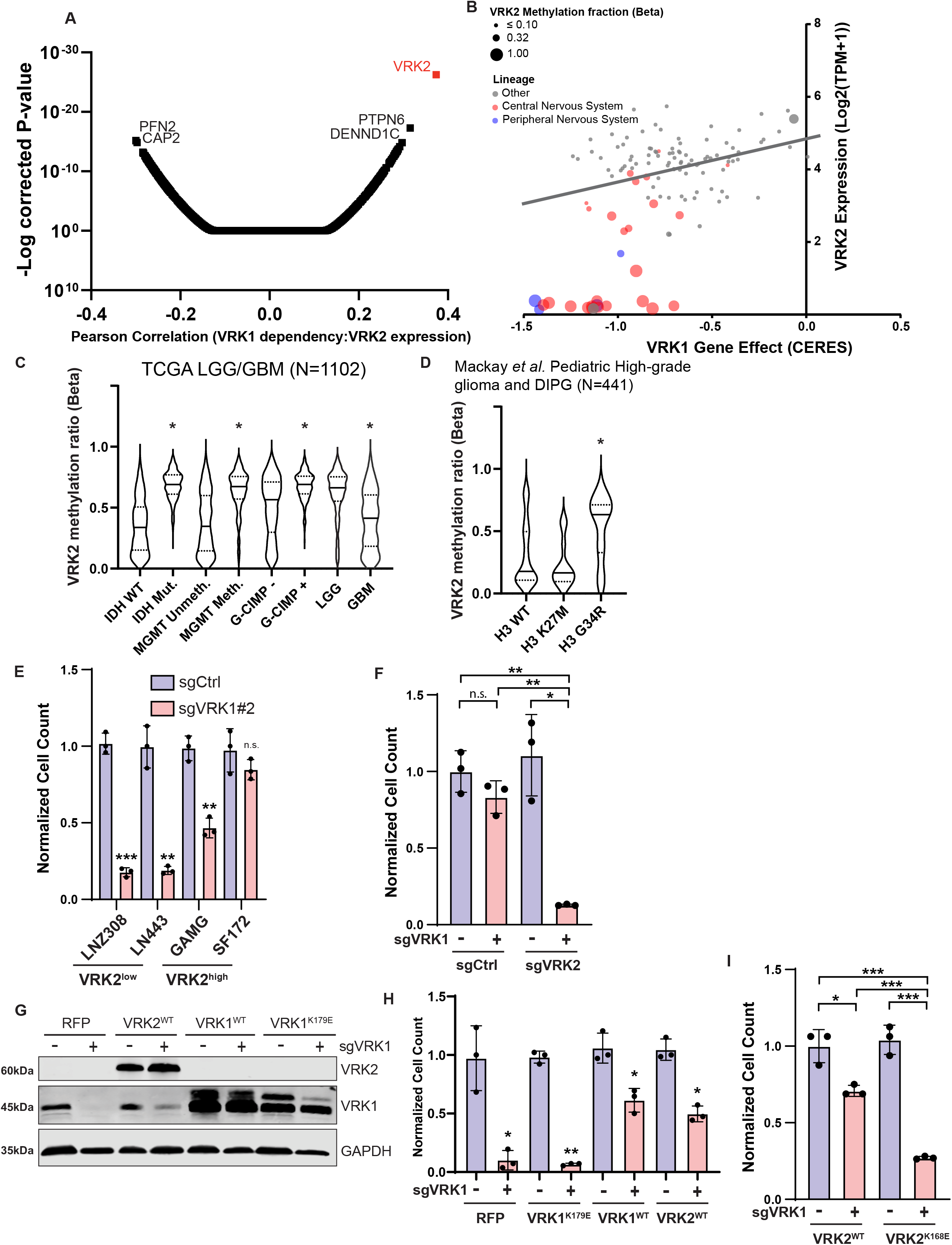
VRK1 dependency is correlated with VRK2 expression. **A**. Whole genome Pearson correlations between gene expression from CCLE (21Q3) and *VRK1* dependency in the Depmap database (21Q3) and adjusted P-values. The topmost correlated genes are annotated. **B**. Scatterplot showing *VRK1* dependency (CERES gene effect) versus *VRK2* expression (log2(TPM + 1)). Extent of *VRK2* promoter methylation is indicated by dot size. Dots are colored according to whether cell lines are in the CNS (red) or PNS (blue) lineages. Only cell lines with DNA methylation data are shown. **C.** *VRK2* promoter methylation status stratified by clinical characteristics across the TCGA GBM-LGG cohort. IDH: Isocitrate dehydrogenase; MGMT: O6-methylguanine-DNA methyltransferase; G-CIMP: Cytosine-phosphate-guanine (CpG) island methylator phenotype; LGG: Low Grade Glioma; GBM: Glioblastoma multiforme. Violin plots with mean (solid line) and 1st and 3rd quartiles (dashed line). **D.** *VRK2* promoter methylation in pediatric high-grade gliomas and DMG with wild-type histone H3 and mutant histone H3 (K27M or G34R). Data from Mackay *et al*., 2017 (39). Violin plots with mean (solid line) and 1st and 3rd quartiles (dashed line). **E.** Cell viability analysis following 14 days KO of *VRK1* in *VRK2*^low^ LNZ308 and LN443 cell lines and *VRK2*^hi^g^h^ GAMG and SF172 cell lines. **F**. Cell viability analysis 14 days following *VRK1* KO in *VRK2*^high^ GBM cell line (SF172), expressing control CRISPR sgRNA or sgRNA targeting *VRK2*. (n=3; mean ± SD). **G**. Immunoblot showing the overexpression of exogenous VRK2^WT^, VRK1^WT^, and kinase-inactive VRK1^K179E^ in NB-1 NB cells with or without *VRK1* KO. **H.** Cell viability analysis for NB-1 cells in panel (G) following 14 days of *VRK1* KO in VRK2^WT^, VRK1^WT^, and kinase-inactive VRK1^K179E^ overexpressing cells. (n=3; mean ± SD). **I.** Effect of VRK2^WT^ or VRK2^K168E^ overexpression on LN443 GBM cell viability following 14 days *VRK1* KO. (n=3; mean ± SD). *p < 0.05, **p < 0.001, ***p < 0.0001; Two-tailed, Student’s T-test for all comparisons.

We next used Celligner, which integrates RNA-sequencing data from TCGA, TREEHOUSE, and TARGET human sequencing studies to identify whether low *VRK2* expression correlates with nervous system cancers for human tumor lineages (10). We found that CNS and PNS tumors exhibit the lowest expression of *VRK2* across all tumor lineages, while *VRK1* expression was similar in CNS/PNS tumors as compared to all other tumor lineages **(Supplementary Fig. S4A, B)**. Within NB, we found that *VRK2* was not differentially expressed among high-MYCN expressing tumors and was slightly elevated in NB tumors enriched for a mesenchymal (MES) gene expression program. However, we note that *VRK2* was still repressed as compared to all other tumor lineages **(Supplementary Fig. S4C).** We also found that in DMG models *VRK2* is not expressed significantly **(Supplementary Fig. S4D)**. These observations suggest that VRK2 is lowly expressed in human cancers of the nervous system, mirroring the cell line (CCLE) data.

Transcriptional repression is enforced through epigenetic regulation, including methylation of CpG dinucleotides at gene promoters (11). Using methylation array data from the TCGA low-grade gliomas and high-grade glioma dataset, we found that *VRK2* promoter methylation occurred more frequently in tumors that exhibit IDH mutations, MGMT methylation, the G-CIMP methylator phenotype or lower grade **(Fig. 2C)**. In a separate dataset of over 1000 pediatric high-grade glioma including DMG, we found *VRK2* promoter methylation in subsets of histone 3 wild-type and H3K27M tumors but most highly associated with the histone H3 G34R mutation **(Fig. 2D)**. We concluded that *VRK2* promoter methylation was an independent predictor for VRK1 dependency.

To experimentally verify the synthetic lethal relationship between *VRK1* and *VRK2*, we focused on a panel of 4 GBM cell lines with heterogeneous expression of *VRK2* **(Supplementary Fig. S5A, B)**. Consistent with our observations in the Cancer Dependency Map, we found greater *VRK1* dependency in the two *VRK2*^low^ cell lines (LNZ308, LN443) than in the two *VRK2*^high^ cell lines (GAMG, SF172) **(Fig. 2E)**.

We then directly tested whether modulation of *VRK2* expression altered the response to *VRK1* KO. To create an isogenic experimental model, we deleted *VRK2* in the *VRK2*^high^ SF172 GBM cell line and then introduced either a control or *VRK1* sgRNA. We found that *VRK2* KO sensitized SF172 to subsequent *VRK1* KO **(Fig. 2F, Supplementary Fig. S5B, C)**. In contrast, ectopic overexpression of wild-type *VRK1*, insensitive to *VRK1* sgRNAs via synonymous mutations, but not kinase-inactive VRK1^K179E^ (12) rescued *VRK1* KO **(Fig. 2G-H, Supplementary Fig. S5D-G)**. Similarly, VRK2 overexpression in *VRK2*^low^ GBM and DMG lines rescued *VRK1* dependency, which required VRK2 kinase activity, as expression of the kinase-inactive VRK2^K168E^ mutant did not rescue *VRK1* KO-induced cell death **(Fig. 2I, Supplementary Fig. S5D-H)**. In summary, *VRK1*-dependent cell lines require VRK1 kinase activity for survival. Furthermore, VRK2 can act as a surrogate kinase for VRK1, providing a mechanistic explanation for the observed VRK1 dependency in CNS and PNS tumors with low VRK2 expression levels.

### Global Phospho-proteomics link VRK1 loss to DNA damage and nuclear membrane substrates

Given the requirement of VRK1 in CNS/PNS tumors, we sought to understand the immediate effects of VRK1 loss. To address this, we designed a degradable VRK1 construct using the dTAG system (13), providing the ability to rapidly deplete exogenous dTAG-VRK1 from cells. Cells were transduced with *VRK1* fused with a C-terminal FKBP12^F36V^ domain (dTAG-VRK1), which can be rapidly degraded with a small molecule (dTAG^V^-1) in a VHL-dependent manner **(Fig. 3A)**. Exogenous expression of dTAG-VRK1 rescued growth defects in the CRISPR KO of endogenous *VRK1* in LN443 (GBM), NB-1 and Kelly (NB) cells, signifying that the fusion protein itself had no effect on canonical VRK1 function **(Fig. 3B-C, Supplementary Fig. S6A)**. However, addition of dTAG^V^-1 and subsequent degradation of dTAG-VRK1 resulted in significantly reduced cell viability in *VRK1* dependent cell lines, establishing a functional system to rigorously examine mechanisms underlying VRK1 dependency **(Fig. 3D, Supplementary Fig. S6A, B, compare Fig. 1D-E)**.

**Figure 3.**
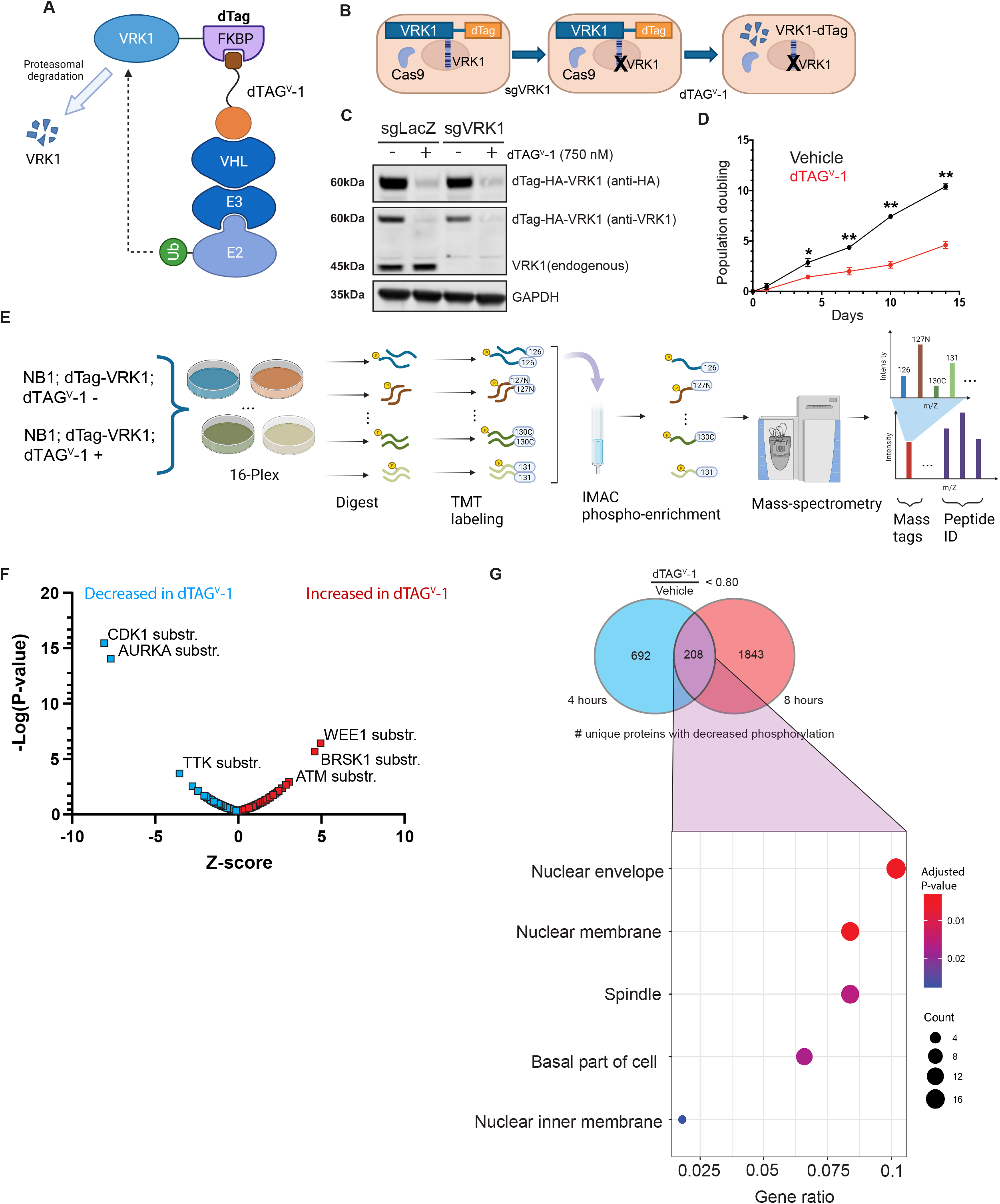
Global phospho-proteomics following acute VRK1 degradation. **A**. Schematic of dTAG-VRK1 degrader system. Based on the FKBP12^F36V^ binding domain, it allows for small-molecule (dTAG^V^-1) mediated recruitment of the VHL ubiquitin ligase complex, targeting exogenous VRK1 for proteasomal degradation. **B**. Schematic of VRK1 degrader experiments. Exogenous dTAG-VRK1 is transduced to rescue CRISPR KO of endogenous *VRK1*. Exogenous dTAG-VRK1 is then under the control of the small molecule degrader (dTAG^V^-1) allowing for acute protein down-regulation. **C.** Immunoblot validation of the dTAG-VRK1 degrader system in NB-1 neuroblastoma cells. Exogenous dTAG-VRK1 was degraded in the presence of dTAG^V^-1. Endogenous *VRK1* was independently targeted with CRISPR KO. sgLacZ is a non-targeting guide control. **D.** Cell viability analysis of dTAG-VRK1-NB-1 cells following addition of either vehicle control or 0.5 μM dTAG^V^-1. Significance at each time point was determined by two-way ANOVA (treatment x time). *p < 0.05, **p < 0.001. **E.** Schematic of the quantitative, global phospho-proteomic experiment. Samples were generated in triplicate at 4h and 8h post dTAG^V^-1 (0.5μM) addition. Following trypsin digestion, peptides are tagged with isobaric, tandem mass tags (TMT), and then combined. Phosphoenrichment was performed using immobilized metal affinity columns (IMAC), and then run on an Orbitrap mass-spectrometer. MS2 spectra offer peptide ID’s as well as sample deconvolution through attached mass tag. **F.** Kinase set enrichment analysis of phosphorylation site dynamics following acute degradation of exogenous VRK1. Kinase substrates of CDK1 and AURKA were significantly down-regulated following degradation (blue), while substrates of WEE1, BRSK1, and ATM were significantly up-regulated (red). **G.** Top panel: Venn diagram showing number of unique proteins with a decrease in phosphorylation (proportion of dTAG^V^-1 / Vehicle <0.8) for at least one amino acid site in dTAG^V^-1 treated samples at 4h and 8h. Bottom panel: Gene set enrichment analysis was performed for the overlap (208 proteins) of downregulated protein phosphorylation at 4 and 8 hours. Dot size indicates the number of genes in each gene ontology category, dot color indicates adjusted P-values, and x-axis is the number of genes in a category from the dataset as compared to the total number of genes.

To identify downstream effectors/pathways of VRK1 kinase, we performed quantitative, phospho-proteomics using the dTAG-VRK1 degrader system **(Fig. 3E)**. dTAG-VRK1-NB-1 cells were treated with dTAG^V^-1 to identify early phosphorylation changes following degradation of VRK1. Following cell lysis, isobaric, tandem mass tagging (TMT) allowed for de-convolution of pooled samples, and relative quantitation among the samples. Phosphorylated peptides were enriched using immobilized metal affinity columns (IMAC) and analyzed by mass spectrometry. We performed kinase set enrichment analysis (KSEA) of phospho-peptide dynamics following either 4 or 8 hours of acute dTAG-VRK1 degradation (14). We found that substrates of cell-cycle and mitotic kinases (CDK1 and AURKA) were down-regulated, while substrates of DNA damage response kinases (ATM and WEE1) were up-regulated **(Fig. 3F)**. A total of 208 phospho-proteins were down-regulated at both the 4h and 8h time-points. Gene set enrichment analysis (GSEA) of these overlapping proteins revealed an enrichment of proteins associated with the nuclear envelope and spindle assembly **(Fig. 3G)**. Notably, members of the inner nuclear membrane, LEM-domain family of proteins, including LEMD3, EMD, and TMPO, showed at least one phosphorylation site that was significantly reduced upon VRK1 degradation **(Supplementary Fig. S7)**. Overall, these observations suggest a critical role of VRK1-regulated pathways in mitosis, nuclear envelope and chromatin homeostasis, as well as DNA damage, in CNS and PNS cell models.

### VRK1 and VRK2 loss leads to post-mitotic nuclear membrane deficits and DNA damage

Phospho-proteomic analysis following VRK1 degradation strongly suggested that VRK1 loss alters the phosphorylation of protein substrates in the nuclear membrane. To visualize nuclear membrane dynamics following VRK1 degradation, we transduced cells with GFP-labelled nuclear lamina-associated proteins: BAF and Emerin. In addition, we also stained cells with anti-LaminB1/2 antibody. Twenty-four hours following degradation of dTAG-VRK1, the nuclear membrane of LN443 GBM cells became misshapen with the formation of lobes and ruffling as well as chromatin bridging between nuclei **(Fig. 4A, B)**. In concordance with reduced cell viability, KO of both *VRK1* and *VRK2* in *VRK2*^high^ cells (SF172) increased irregular nuclei compared to individual kinase KO alone **(Fig. 4C, Supplementary Fig. S8A)**.

**Figure 4.**
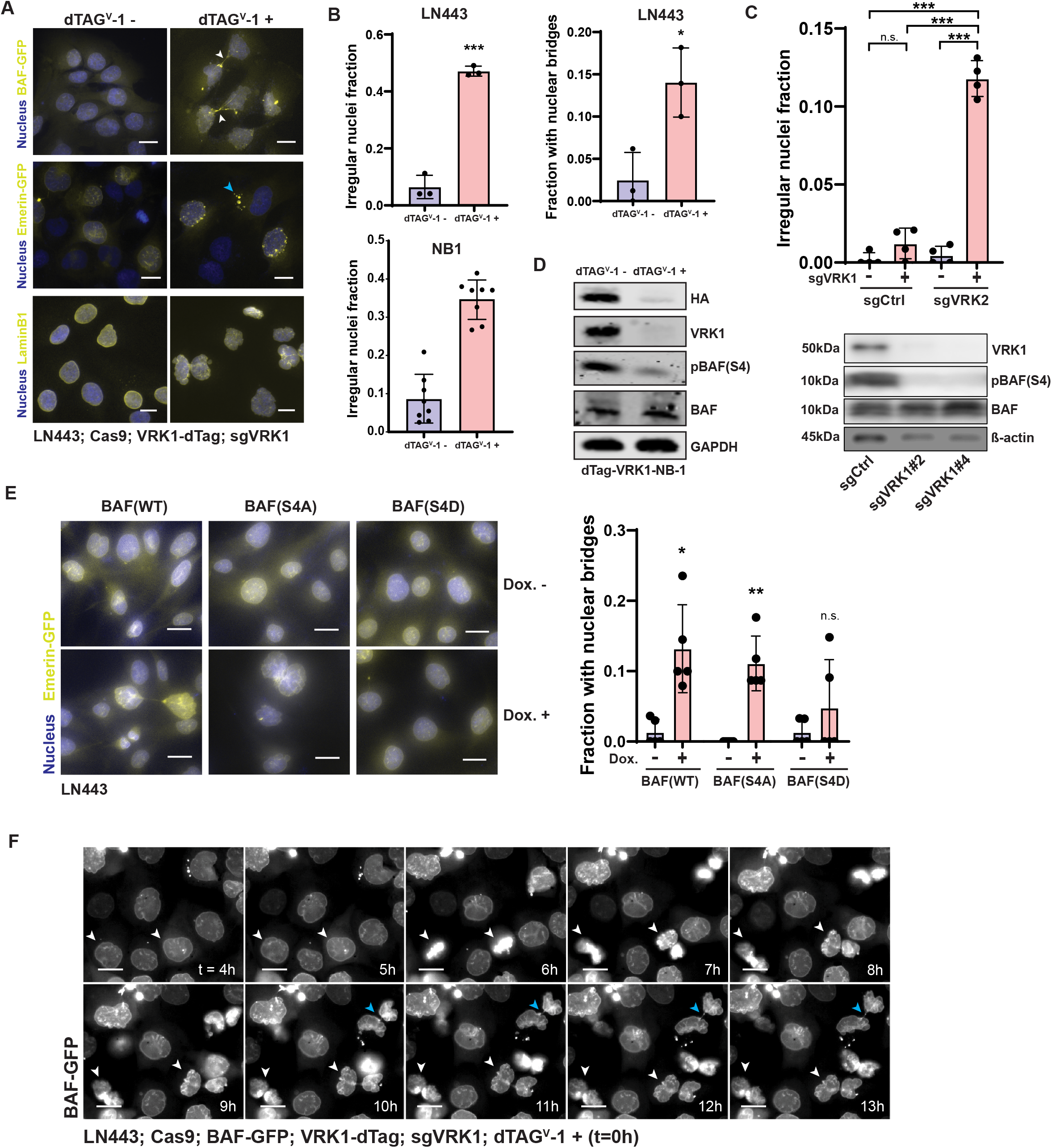
VRK1 loss is associated with nuclear envelope malformation. **A**. Nuclear membrane morphology in the LN443 GBM cell line following exogenous VRK1 degradation by dTAG^V^-1 (0.5 μM) after 1 day. Nuclear membrane visualized by GFP-tagged BAF and Emerin as well as immuno-fluorescent staining for LaminB1. White arrows point to nuclear bridges. Blue arrow points to micro-nuclei. **B**. Top, left: Quantitation of the fraction of irregular nuclei, by LaminB1 staining, following VRK1 degradation as seen in **Fig. 4A** (n=3 fields of >50 cells each; mean ± SD). Top, right: Quantitation of the fraction of nuclear bridges following VRK1 degradation as seen in **Fig. 4A** (n=3 fields of >50 cells each; mean ± SD). Bottom, left: Quantitation of the fraction of irregular nuclei following VRK1 degradation in the NB-1 NB cell line expressing GFP-BAF seen in **Supplementary Fig. S8D** (n=8 fields of >50 cells each; mean ± SD). **C.** Quantitation of irregular nuclei, by LaminB1 staining, following KO of both *VRK1* and *VRK2* in SF172 as seen in seen **Supplementary Fig. S8A.** (n=4 fields of >50 cells each; mean ± SD). **D.** Immunoblot of phosphorylated BAF (S4) and total BAF following dTAG^V^-1 treatment in dTAG-VRK1-NB-1 cells (left panel) or KO of *VRK1* with two independent sgRNA in BT869Luci DMG neurospheres (right panel). Representative of 2 independent experiments. pBAF and total BAF were probed in two separate blots of same lysates. **E.** Left panel: Nuclear envelope morphology as seen by GFP-tagged Emerin following 0.5 μM doxycycline-induced expression of BAF mutants in LN443 GBM cell line after 3 days: wild-type (WT), S4A (non-phosphorylatable), S4D (phospho-mimetic). Right panel: Quantitation of nuclear bridging phenotype in LN443 cell lines expressing BAF mutants (n=3; mean ± SD). **F.** Time-lapse of live-cell experiment showing nuclear envelope morphology (GFP-tagged BAF) following VRK1 degradation in LN443 (dTAG^V^-1 addition at t=0h). White arrows point to cells undergoing mitosis. Blue arrows point to chromatin bridges. Representative of 2 independent experiments. Scale bars: 20 μm. *p < 0.05, **p < 0.001, ***p < 0.0001; Two-tailed, Student’s T-test for all comparisons.

The nuclear envelope protein BAF serves to tether chromatin to proteins in the inner nuclear membrane and is a substrate of both VRK1 and VRK2 individually on the Serine-4 (S4) residue (15,16). During mitosis, phosphorylation of BAF on Serine-4, and subsequent nuclear lamina-DNA un-tethering, is required for mitotic chromosome segregation, as well as post-mitotic nuclear envelope re-assembly (16). BAF(S4) phosphorylation was not detectable in our phospho-proteomic analysis, however, the observed reduced phosphorylation of LEM-domain proteins that bind to BAF **(Supplementary Fig. S7)** led us to hypothesize that the altered nuclear envelope dynamics observed upon *VRK1* KO was due to decreased phosphorylation of BAF. Indeed, dTAG^V^-1-mediated degradation of exogenous VRK1 in dTAG-VRK1-NB-1 cells and CRISPR KO of *VRK1* in the *VRK2*^low^ DMG neurosphere models BT869Luci and SU-DIPGXIIIP*Luci strongly decreased levels of phosphorylated BAF (S4) but not total BAF **(Fig. 4D, Supplementary Fig. S8B)**. We tested whether ectopic overexpression of the non-phosphorylatable mutant BAF^S4A^ would mimic *VRK1/2* loss, and indeed, following doxycycline-induced expression, we observed similar nuclear bridges and distorted nuclear envelope morphology **(Fig. 4E, Supplementary Fig. S8C)**. We also found that doxycycline-induced ectopic overexpression of BAF^WT^ resulted in the same phenotype, perhaps by saturating the phosphorylation capacity of VRK1/2, leading to a shift in the pool of BAF towards its unphosphorylated form **(Fig. 4E, Supplementary Fig. S8C)**. In contrast, overexpression of the phospho-mimetic mutant BAF^S4D^ had no effect on nuclear morphology **(Fig. 4E, Supplementary Fig. S8C)**, suggesting that the Serine-4 phosphorylation site plays a crucial role in VRK1 kinase dependency. Using live-cell imaging, we followed BAF dynamics after dTAG-VRK1 degradation in LN443 GBM cells **(Fig. 4F)**. We observed the same nuclear envelope ruffling and bridging in cells immediately following mitosis. We also found these same phenotypes in NB-1 NB cells, where nuclear membrane ruffling predominated **(Supplementary Fig. S8D)**. Taken together, these findings indicate that BAF Serine-4 phosphorylation by VRK1 is essential for CNS and PNS tumor cells to maintain the integrity of nuclear envelope structure and function.

In addition to observing altered protein phosphorylation at a number of proteins in the nuclear envelope, we also noted an enrichment for substrates of the DNA damage pathways (i.e. substrates of ATM and WEE1) **(Fig. 3F)**. Therefore, we performed imaging of DNA damage response foci. At 7 days following KO of *VRK1*, we found an increased number of phospho-H2AX foci (S139), phospho-ATR (S428), and phospho-DNAPK (S2056), representing induction of both non-homologous end-joining and homologous recombination pathways of DNA double-strand break repair **(Fig. 5A)**. Corroborating potentiated apoptosis induction, concomitant KO of *VRK1* and *VRK2* increased DNA damage foci (phospho-H2AX) in *VRK2*^high^ GBM cells **(Fig. 5B)** and in two NB cell lines after degradation of dTAG-VRK1 **(Fig. 5C)**.

**Figure 5.**
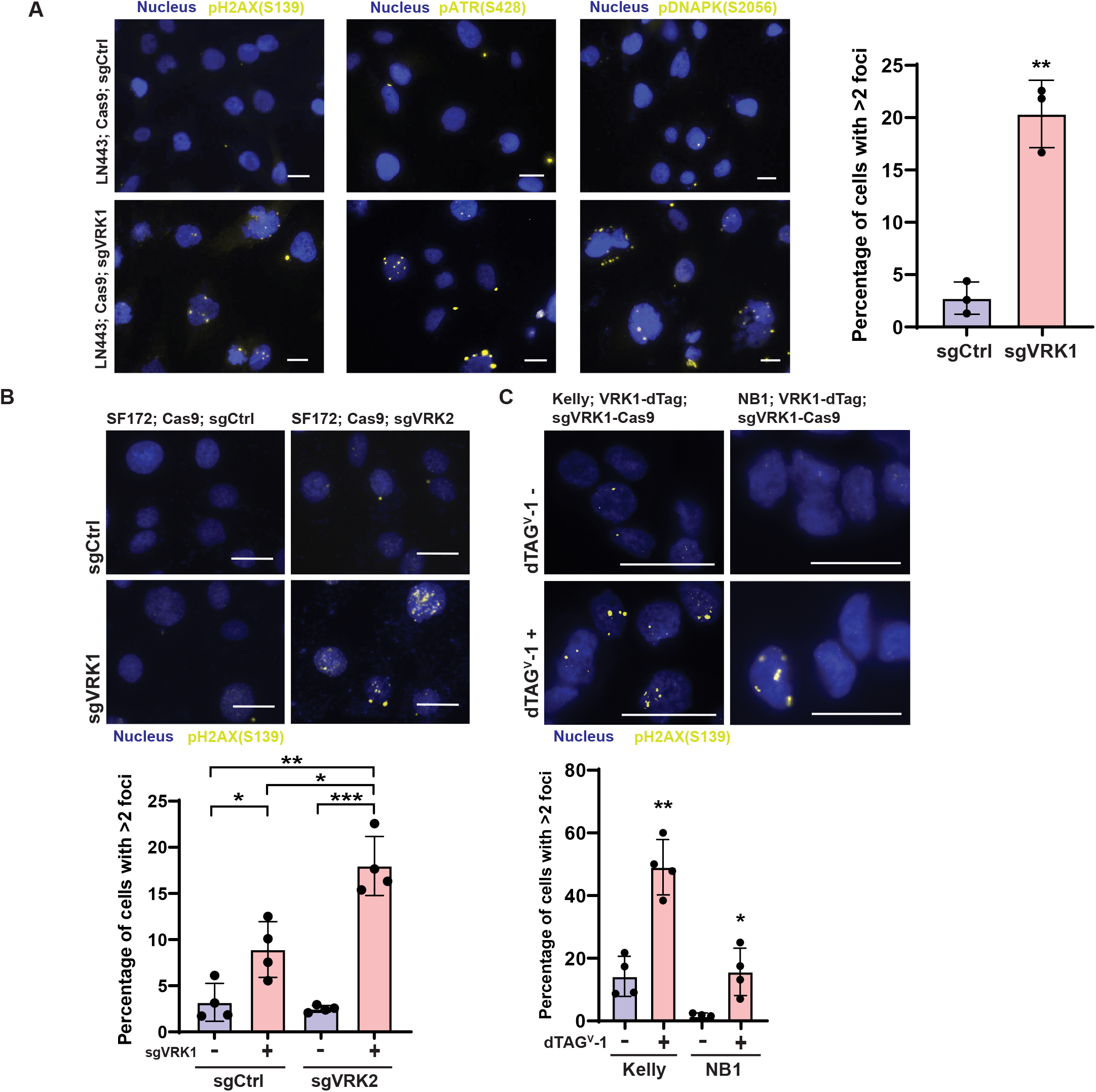
VRK1 loss results in DNA damage. **A**. Left panel: Nuclear foci of a panel of DNA damage markers (phospho-H2AX(S139), phospho-ATR(S428), phospho-DNAPK(S2056)) following KO of *VRK1* in LN443 GBM cells for 7 days. Right panel: Quantitation of percent of cells with >2 phospho-H2AX foci following *VRK1* KO (n=3 fields of >50 cells each; mean ± SD). **B**. Top panel: Phospho-H2AX foci following 7 days double KO combinations of sgCtrl/sgCtrl, sgCtrl/sgVRK1, sgCtrl/sgVRK2, and sgVRK1/sgVRK2. Bottom panel: Quantitation of percent of cells with >2 phospho-H2AX foci following these double-KO combinations (n=4 fields of >50 cells each; mean ± SD). **C.** Top panel: Phospho-H2AX foci following VRK1 degradation with 0.5μM dTAG^V^-1 in both Kelly and NB-1 neuroblastoma cell lines. Bottom panel: Quantitation of percent of cells with >2 phospho-H2AX foci following dTAG^V^-1 addition (n=4 fields of >30 cells each; mean ± SD). Scale bars: 20 μm. *p < 0.05, **p < 0.001, ***p < 0.0001; Two-tailed, Student’s T-test for all comparisons.

### VRK1 is a dependency in tumor models in vivo

To evaluate *VRK1* dependency *in vivo*, we utilized a tamoxifen-inducible CRISPR-Cas9 system (17). Plasmids expressing Cas9, Cre-ERT2, and the pLenti_Switch-ON guide plasmid targeting *VRK1* were transduced into LN443 or SF295 GBM cell lines. The “Switch-ON” plasmid has CRISPR guide expression suppressed with a LoxP-STOP-LoxP site. Upon tamoxifen treatment, Cre recombinase is induced which removes the transcriptional stop, and allows expression of the guide RNA. We first validated *VRK1* KO efficiency *in vitro* and also observed decreased viability after VRK1 depletion **(Fig. 6A).** This cell line (SF295; Cas9; CreERT2; pLenti_Switch-ON_sgVRK1) was subsequently injected into flanks of NSG mice **(Fig. 6B)**. Once the xenografts reached ~200 mm^3^, *VRK1* KO in tumor cells was induced by intraperitoneal administration of tamoxifen. *VRK1* KO resulted in virtually complete and durable tumor remission in all mice (n=10 tumors) 10-20 days post tamoxifen treatment, whereas tumors in vehicle-treated controls continued exponential growth **(Fig. 6C)**. Animals were euthanized when their tumor exceeded 500 mm^3^ or 40 days following tamoxifen administration. For a subset, we harvested tumors 7 days following treatment with tamoxifen or vehicle control, stained for phospho-H2AX (S139) and found evidence of increased DNA damage in tumors with *VRK1* depletion from tamoxifen treatment **(Fig. 6D)**. These observations confirm that *VRK1* depletion leads to tumor regression *in vivo*, suggesting that VRK1 is a potential therapeutic target in *VRK2* promoter-methylated adult and pediatric gliomas and neuroblastomas.

**Figure 6.**
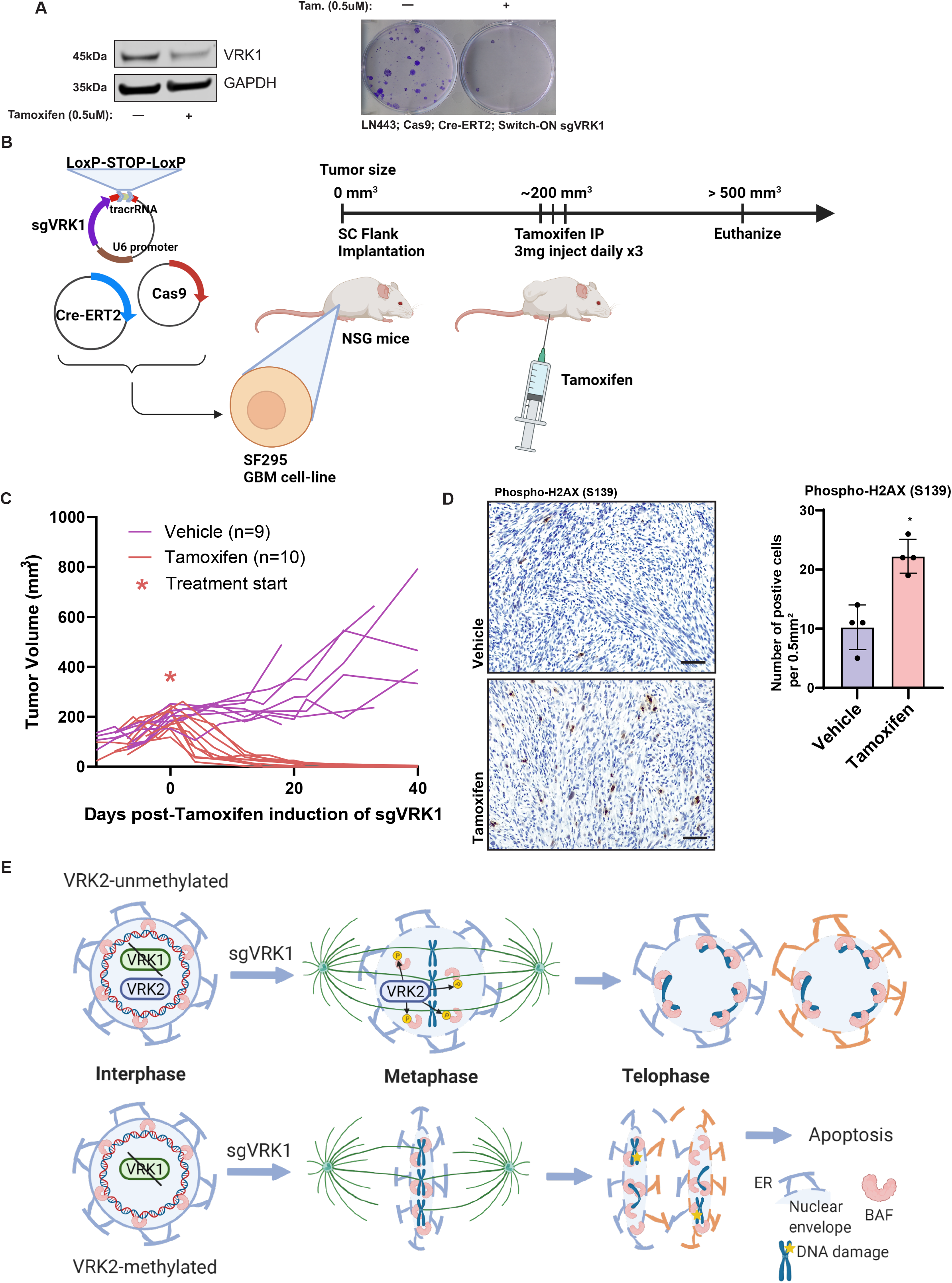
VRK1 is a dependency *in vivo*. **A**. Left panel: Immunoblot of VRK1 following tamoxifen-induced expression of sgVRK1 in LN443 cells. Right panel: Clonogenic assay in LN443 cells 14 days following tamoxifen-induced KO of *VRK1*. **B**. Schematic of the *in vivo* xenograft experiment. The SF295 GBM cell line was transduced with Cas9, Cre-ERT2, and “Switch-ON” guide plasmids and implanted in NSG mouse flanks. When the tumors reached a pre-specified size (200 mm^3^), the mice were treated with tamoxifen. When the tumor size reached ~500 mm^3^ or 40 days following treatment, the mice were euthanized. **C**. Tumor volume measurements over time of the flank xenografts. * represents injection of tamoxifen or corn oil vehicle control. **D**. Left panel: representative H&E sections of tumors taken from xenografted mice, 7 days following treatment with tamoxifen or vehicle control (Scale bar: 50μm). Sections were stained with antibody against phospho-H2AX. Right panel: quantitation of number of phospho-H2AX positive cells per 0.5 mm^2^ in flank xenografts following tamoxifen or vehicle treatment (n=4 fields; mean ± SD) (*p < 0.05; Two-tailed, Student’s T-test). **E.** Schematic showing the proposed mechanism of synthetic lethality between *VRK1* and *VRK2*. In *VRK2* unmethylated tumors (top), VRK2 compensates for loss of *VRK1* in the phosphorylation of BAF during mitosis. In *VRK2*^low^ tumors (bottom), loss of *VRK1* leads to retention of BAF during mitosis and the continued association of the nuclear envelope with chromatin. This leads to impaired chromosomal segregation and DNA damage, as well as nuclear bridging.

## Discussion

Synthetic lethal interactions are a potential source of new biomarker-linked targeted cancer therapy. Specifically, synthetic lethal interactions may involve tumor-specific down-regulation of a gene or pathway, resulting in sensitivity to inhibition of another gene or pathway. The success of PARP inhibitors in multiple cancers with homologous recombination pathway deficiency provides evidence that this approach can lead to clinical benefit (18,19). Specifically, BRCA1/2 mutations in breast cancer result in dependency on the non-homologous end-joining DNA repair pathway that is exploited by PARP inhibitors like Olaparib (20).

Gene paralogs are potentially promising sources of synthetic lethal interactions as they usually exhibit strong sequence homology and functional redundancy. For example, alpha-enolase (*ENO1*)-deleted GBMs are sensitive to KO of its paralog gamma-enolase (*ENO2*), blocking glycolysis (21). Loss-of-function *ARID1A*-mutant cancers are sensitive to *ARID1B* KO, causing destabilization of the SWI/SNF chromatin remodeling complex (22). Synthetic lethality in the context of paralogs can occur by epigenetic mechanisms as well. For example, in *NXT2*-methylated NB cell lines, NXT1 is required to facilitate stability of the essential RNA-exporting protein NXF1 (23). Targeting paralogs holds the promise of an increased therapeutic ratio as one interaction partner may be a silenced tumor suppressor or may be co-silenced with other tumor suppressors but not affected in normal tissues. Here, we discovered that tumors with low VRK2 expression are dependent on its paralog, VRK1. IDH-mutant gliomas, with their hyper-methylated phenotype, also exhibit high *VRK2* gene methylation. In fact, *VRK2* promoter methylation is highly enriched in tumors of the CNS and PNS lineages. During development, differential gene methylation is involved in neuronal cell-fate determination, neuronal plasticity, and memory formation (24). Such lineage-specific, differential methylation may lead to other synthetic lethal vulnerabilities in cancer.

The VRK family of atypical Serine-Threonine kinases was initially discovered for their homology with vaccinia virus B1 kinase, which is required for viral replication (25). The family branches early from the kinase evolutionary tree and consists of the functional kinases VRK1 and VRK2 and a pseudokinase VRK3 (12). Clinically, VRK1 expression has been associated with high grade and poor prognosis in patients with glioma (26), whereas *VRK2* expression is correlated with improved survival in high-grade astrocytoma (27). In the physiological context, VRK1 localizes to the nucleus where it is thought to phosphorylate substrates involved in DNA damage response (e.g. histone H2AX) and mitosis (e.g. BAF) (12). Previous work showed that VRK1 is the primary kinase that phosphorylates BAF during mitosis (16). BAF phosphorylation removes its association with chromatin and LEM-domain containing proteins of the nuclear envelope, such as Emerin. Like VRK1, VRK2 is known to phosphorylate BAF, modulating its association with the nuclear membrane in mitosis (15). Unlike VRK1, which localizes to the nucleoplasm, VRK2 associates with A-type Lamins of the nuclear envelope. Birendra *et al*. hypothesized that VRK1 may modulate BAF phosphorylation in the nucleoplasm, while VRK2 modulates BAF at the nuclear envelope (15). This difference in localization may explain the only partial rescue of VRK1 loss by VRK2 that we observed (**Fig. 2H, I**).

Together, our observations are consistent with a synthetic lethal interaction of *VRK1* and *VRK2* **(Fig. 6E)**. In *VRK2*^high^ tumors where the *VRK2* promoter is unmethylated, both VRK1 and VRK2 may phosphorylate BAF during mitosis to mediate nuclear envelope dis-assembly. However, in *VRK2*^low^ tumors, loss of *VRK1* prevents BAF phosphorylation during mitosis. Thus, our data suggest that VRK1 depletion results in retained association of nuclear envelope fragments with mitotic chromosomes, leading to aberrant nuclear envelope re-assembly, nuclear bridging between daughter cells, and ultimately to DNA damage and apoptotic cell death.

Small molecule kinase inhibitors have been investigated for their potential differential effect on VRK1 versus VRK2 activity (28,29). Vázquez-Cedeira *et al*. noted that, based on amino-acid sequence and protein structural differences from other kinases, both VRK1 and VRK2 are predicted to have low promiscuity and be relatively insensitive to extant kinase inhibitors (28). They further showed that in a small-molecule library screen of 20 kinase inhibitors, few molecules decreased VRK1 or VRK2 kinase activity even at high concentrations (100 μM). The compounds that did inhibit kinase activity did so with ATP concentrations three orders of magnitude lower than intracellular levels, which the authors noted may limit *in vivo* use. Recently, a small molecule, based on an aminopyridine scaffold, was developed that showed potent activity against VRK1 *in vitro* (IC50 = 150 nM) (29). However, this compound did not significantly decrease viability in cell culture (29). Potent kinase inhibitors that show differential effect against VRK1 versus VRK2 do not yet exist. A degrader strategy, as modeled in this current study, may represent an alternate approach to targeting VRK1 as a growing number of small-molecule degraders (e.g. PROTACs, molecular glues, etc.) targeting specific proteins are undergoing clinical trials in diverse cancers (30).

For VRK1 inhibition to be a viable therapy option, a significant therapeutic ratio is required where normal tissues are spared while cancer cells are targeted. The existence of human genetic variants and mouse transgenic models allow for a glimpse of potential on-target toxicities. A rare germ-line mutation in VRK1 (R358X) results in lack of VRK1 protein production, and manifests in pediatric patients as spinal muscular atrophy with pontocerebellar hypoplasia (SMA-PCH) (31). Although *VRK2* is expressed in most tissues, it has low expression in normal brain tissue, especially the cerebellum, which may explain the CNS phenotype of mutant VRK1 **(Supplementary Fig. S3)**. Partial KO of *Vrk1* by gene-trapping resulted in a slight reduction in brain size, mild motor dysfunction, and male infertility in mice (32,33). These findings suggest that side effects of VRK1 inhibition may be tolerated in adults.

In summary, by integrating genome-wide, loss-of-function genetic screens with RNA-sequencing and DNA methylation, we identified VRK1 as a selective vulnerability in CNS and PNS cancers with low *VRK2* expression. Taken together, these studies suggest that targeting VRK1 in cancers that harbor promoter-methylated *VRK2* is a potential therapeutic strategy.

## ACKNOWLEDGMENTS

We thank members of the Hahn Lab, Stegmaier Lab, and Filbin Lab, and Nilay Sethi for useful discussions and technical assistance. This work was supported by NIH U01 CA176058 (WCH), NIH R03 TR 003343 (WCH and JS), NIH R35 CA210030 (KS), NIH 1P01 CA217959 (KS), NCI R25 CA174650 (BS), NCI F32 CA243290 (JK), and the Erwin Schrödinger Fellowship of the Austrian Science Fund, Vienna, Austria (#J-4311, B.E.). M.G.F. holds a Career Award for Medical Scientist from the Burroughs Wellcome Fund, the Distinguished Scientist Award from the Sontag Foundation and the A-Award from the Alex’s Lemonade Stand Foundation. NWM was supported by the National Cancer Institute under a Ruth L. Kirschstein National Research Service Award (F32 CA261035) and the Dana-Farber Cancer Institute Ungerer Fellowship award. E.P. holds a Fellowship funded by the Michael Mosier Defeat DIPG Foundation, The ChadTough Foundation, and SoSo Strong Pediatric Brain Tumor Foundation.

We thank Lai Ding and the NeuroTechnology Studio at Brigham and Women’s Hospital for providing live-cell imaging instrument access and consultation on data acquisition. We also thank Mark Jedrychowski, Julian Mintseris, and the Thermo Fisher Scientific Center for Multiplexed Proteomics at Harvard Medical School (http://tcmp.hms.edu) for mass spectrometry data acquisition and analysis.

## AUTHORS’ CONTRIBUTIONS

**J. So:** Conceptualization, data curation, formal analysis, investigation, writing–original draft. **N. Mabe:** Conceptualization, data curation, formal analysis, investigation, writing–original draft. **B. Englinger:** Conceptualization, data curation, formal analysis, investigation, writing–original draft. **S. Moyer:** Formal analysis, investigation. **B. Shim:** Formal analysis, investigation. **M.C. Trissal:** Investigation. **J. Marques:** Investigation. **J. Kwon:** Investigation. **E. Panditharatna:** Investigation. **D. Jeong:** Investigation. **D. Mayhew:** Investigation. **J. Hwang:** Investigation. **M.G. Filbin:** Conceptualization, resources, supervision, funding, writing–original draft. **K. Stegmaier:** Conceptualization, resources, supervision, funding, writing–original draft. **W.C. Hahn:** Conceptualization, resources, supervision, funding, writing–original draft.

## Materials and Methods

### EXPERIMENTAL MODEL AND SUBJECT DETAILS

#### Cell Culture

Neuroblastoma (NB-1, Kelly) and GBM (LN443, SF172, GAMG, LNZ308) cell lines were collected from the Cancer Cell Line Encyclopedia and Cancer Dependency Map projects and obtained from the Broad Institute. The cell lines that express pLX_311-Cas9 were generated by Project Achilles (https://depmap.org/portal/achilles)(20). SK-N-BE(2)C were purchased from ATCC. LAN-1 was kindly gifted by Rani George at DFCI. SK-N-BE(2)C, LAN-1, and GBM cell lines were grown in 10% DMEM supplemented with glutamine, penicillin and streptomycin, and incubated at 37°C in 5% CO2. Kelly and NB-1 were grown in RPMI-1640 supplemented with 10% FBS and glutamine, penicillin, and streptomycin, and incubated at 37°C in 5% CO2. Cell lines identities were validated by STR profiling and tested negative for mycoplasma with MycoAlert Mycoplasma Detection Kit (Lonza, Cat#LT07-418) prior to experimental use.

#### Neurosphere Culture

Patient-derived H3K27M and H3WT-glioma neurosphere lines were established at Boston Children’s Hospital (BT869/BT869Luci), Stanford University (SU-DIPGXIIILuci, SU-DIPGXIIP*Luci, SU-DIPGXXV, SU-pcGBM2, SU-DIPG48), and Hospital Sant Joan de Deu Barcelona (HSJD-DIPG007, HSJD-GBM001) as previously described (34–36). H3K27M-glioma cells were grown as neurospheres in tumor stem media (TSM) base (36) supplemented with B27 minus vitamin A (Thermo Fisher Scientific), human growth factors (EGF, FGF, PDGF-AA, PDGF-BB [Shenandoah Biotechnology]) and heparin (Stemcell Technologies) in ultra-low attachment flasks. Indicated cell models expressing luciferase were generated as described in (37). Neurosphere cultures were dissociated for passaging using Accutase cell detachment solution (Stemcell Technologies) for 3-5 minutes at 37°C. All neurosphere models were authenticated by high resolution short tandem repeat (STR) profiling (Molecular Diagnostics Core, Dana-Farber Cancer Institute). Whole exome or whole genome sequencing was conducted on neurosphere models to obtain copy number alterations.

### METHOD DETAILS

#### Public Data Sets

Log2(TPM) + 1 RNA-sequencing, CERES gene dependency scores, and DNA methylation array data were downloaded from the Dependency Map portal (https://depmap.org/portal/, CCLE expression: 21Q3). Density plots displaying the distribution of CERES scores per tumor lineage were generated with ggridges software in R (v4.0.3). Projection of *VRK2* or *VRK1* expression for tumor lineages from TCGA/TARGET/TREEHOUSE tumor datasets was generated using UMAP projection plots available in the Celligner alignment portal (available from: depmap.org/portal/celligner). TCGA LGG and GBM gene expression and clinical data downloaded from cbioportal.org. Methylation data for pediatric high-grade gliomas from Mackay *et al*. downloaded from ArrayExpress (https://www.ebi.ac.uk/arrayexpress/experiments/E-MTAB-5528/) (39).

#### Other Data Analysis and Statistics

For statistical tests of significance, the statistical test and P-value are described in the respective figure legends. All t tests are two-sided unless otherwise indicated. A P value of 0.05 was used as the cutoff for significance unless otherwise indicated. These values were calculated in GraphPad Prism (version 9.3.0 for Windows, GraphPad Software, San Diego, California USA, www.graphpad.com) or R version 4.0.2 and Rstudio version 1.2.5042. Error bars represent SD unless otherwise indicated. All duplicate measures were taken from distinct samples rather than repeated measures of the same sample.

#### Lentiviral production

Lentiviral production was conducted using HEK293T cells, as described on the Broad Institute Genetic Perturbation Platform (GPP) web portal: (https://portals.broadinstitute.org/gpp/public/). Briefly, high-titer lentivirus was produced by transfection of HEK293T cells with the lentiviral vector, psPAX2 (Addgene #12260) and vsvg (Addgene #8454) with Lipofectamine 2000 (Life Technologies Cat#11668027). Viral supernatant was collected 48 hours after transfection and filtered with 0.2 micron filter. Cells were transduced with virus in the presence of 5 μg / mL polybrene and selected with blasticidin (5 μg / mL) or puromycin (1 μg / mL) according to appropriate selection agent. dTAG-HA-VRK1-expressing cell lines were derived by first expressing stable dTAG-HA-VRK1 prior to infection with sgVRK1#2.

#### Single guide RNAs (sgRNAs)

The single guide RNA (sgRNA) sequences used for the validation experiments were designed using the web-based program (CRISPick) provided by the Broad Institute GPP: (https://portals.broadinstitute.org/gppx/crispick/public). For the CRISPR-mediated gene KO (KO), annealed oligonucleotides carrying the sgRNA target sequence as well as the cloning adapters were inserted into a guide RNA-expressing vector that also expresses a puromycin-resistance gene (pXPR_003, Broad Institute GPP), the vector expressing the hygromycin-resistance gene (pXPR_016, Broad Institute GPP), or guide vectors with GFP or mCherry coexpression (LCV2_EGFP or LCV2_mCherry). LCV2_EGFP and LCV2_mCherry were gifts from Jason Moffat (Addgene plasmid #155098 and #155096) (40). The targeting sequences for the individual sgRNAs were:

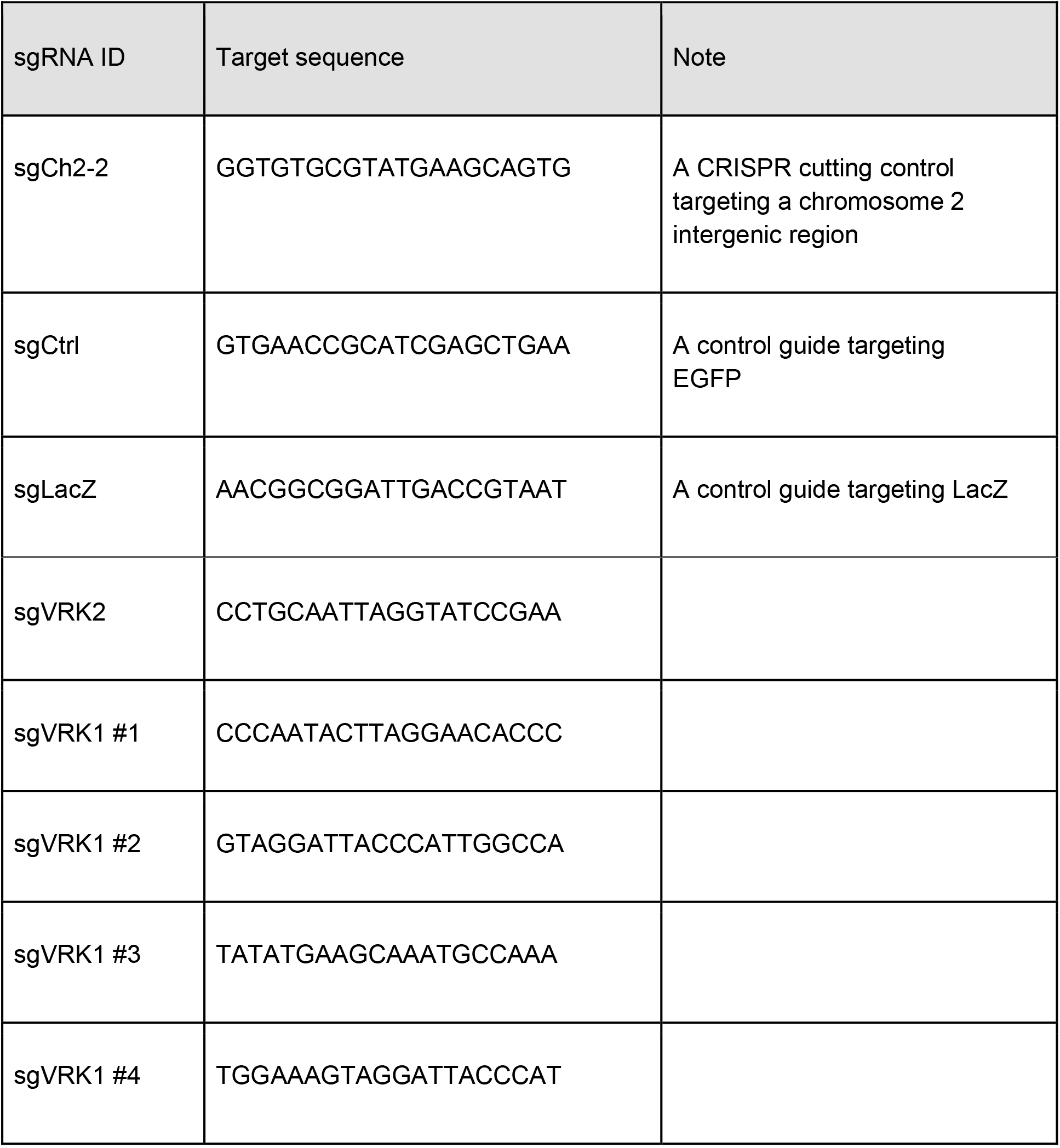

For tamoxifen inducible sgRNA expression, we utilized the CRISPR-Switch system as described by Chylinski *et al*. (17). Guides were cloned into the vector pLenti_Switch-ON which was a gift from Ulrich Elling.

#### Open Reading Frame (ORF) Constructs

Codon-optimized, sgRNA-resistant DNA fragments encoding VRK1^WT^, VRK1^K179E^, VRK2^WT^ were purchased from gBlock (IDT) and cloned into pDONR-221 via BP gateway cloning. VRK2^K168E^ was generated through the QuickChange II site-directed mutagenesis kit (Agilent Technologies) using the primer: 5’-GAATATGTTCATGGTGATATAGAAGCAGCAAATCTAC-3’. BAF^WT^ and its mutants (S4A and S4D) were synthesized with Gateway-compatible AttP flanking sites (IDT) and also cloned into pDONR-221. Entry clone pENTR/D_creERt2 was a gift from Leonard Zon (Addgene plasmid # 27321) (41). VRK1^WT^ was further cloned into PLX305(C-TAG) (Addgene #91798) and VRK1^WT^, VRK1^K179E^, VRK2^WT^, VRK2^K168E^, and creERt2 were further cloned into PLX307 (Addgene #41392) via LR Gateway cloning (LR clonase II enzyme mix, Thermo-Fisher Scientific, Cat# 11791-100). BAF^WT^, BAF^S4A^, and BAF^S4D^ were cloned into Doxycycline-inducible expression vector PLXI403 (Addgene #41395). Cells were transduced with virus in the presence of 5 μg / mL polybrene and selected with blasticidin (5 μg / mL) or puromycin (1 μg / mL) according to appropriate selection agent. dTAG-HA-VRK1-expressing cell lines were derived by first expressing stable dTAG-HA-VRK1 prior to infection with sgVRK1#2.

Further information and requests for reagents should be directed and fulfilled by the lead contact William Hahn. Plasmids for the C-terminus, dTAG-VRK1, sgRNAs#1,#2, and #4, and cDNAs for VRK1^WT^, VRK1^K179E^, VRK2^WT^, VRK2^K168E^, BAF^WT^, BAF^S4A^, and BAF^S4D^ will be made available on Addgene.

#### Cell Proliferation Assay

The viability effect of *VRK1* KO in GBM and DMG cell lines was determined by the clonogenic cell proliferation assay. Briefly, cells were transduced with guide RNA sgCtrl or sgVRK1. Following 1 wk under selection, 0.5-1×10^4^ cells per well were seeded in 6-well Falcon plates (Fisher Scientific, Cat #087721B) in triplicate. Media was changed every 5-7 days. After 7-10 days, cell numbers were counted using the Vi-Cell automated cell counter (Beckman Coulter, Cat # 731196).

For viability effect in neuroblastoma cell lines, 5 × 10^5^ cells of either sgChr2, sgVRK1#1, sgVRK1#2, or sgVRK1#4 were plated onto 6-cm dishes. For dTAG-VRK1 cells, 5 × 10^5^ cells were plated and attached 16 hours prior to incubation with either DMSO vehicle or 1 μM dTAG^V^-1. After 2-3 days, cells were detached, counted, and the number of doublings relative to the prior timepoint were calculated. Groups were replated at 5 × 10^5^ cells per group, and the same steps were repeated every 2-3 days for a total of 14 days. For days in which fewer than 5 × 10^5^ cells were counted, then all the cells were plated. Population doublings were calculated by the total cells compared to the number of seeded cells. Values were added to the previous time point, starting at 0 for day 0. dTAG-VRK1 cells remained in vehicle or 1 μM dTAG^V^-1 for the entirety of the 14 days.

For crystal-violet staining, cells were plated in 6-well plates and stained/fixed with 2.5mg/mL solution of crystal-violet (Sigma, Cat #C3886) in 20% methanol.

#### Cell-cycle and Apoptosis Assay by Flow Cytometry

Cells were harvested, washed, and then fixed in ice-cold 70% ethanol and then re-suspended in stain buffer containing propidium iodide and RNase (BD, Cat # 550825). Apoptosis was assessed using annexin V and propidium iodide staining according to the manufacturer’s instructions (Invitrogen, Cat # 88-8005-74). Samples were analyzed on a BD LSR-II flow cytometer. Data analysis was completed using the cell-cycle analysis package in FlowJo ver.10.8.0 (Treestar).

#### Western Blot

Cell pellets were lysed with CST lysis buffer (Cat #9803) that was supplemented with phosphatase (Roche, Cat #04906845001) and protease inhibitors (Roche, Cat #11836170001) and diluted to 1 μg / μL in sample buffer.

Approximately 35 μg of whole-cell lysate protein was loaded into wells and resolved in 4-12% acrylamide gradient gels. Whole cell lysates were run with MOPS running buffer solution for high molecular weight proteins and MES running buffer solution for low molecular weight proteins. Acrylamide gels were wet-transferred onto nitrocellulose or PVDF membranes for at least 90 minutes. Primary antibodies found against VRK1 (CST, Cat #3307), VRK2 (Life Technologies, Cat #MA527456), BAF (Life Technologies, Cat #MA534813), GAPDH (CST, Cat #2118), HA (CST, Cat #2367), or β-actin (CST, Cat#3700) were diluted in 3% BSA in TBS-T and incubated overnight at 4 degrees Celsius. Rabbit polyclonal anti-phospho-BAF antibody was a gift from Dr. Robert Craigie (National Institute of Health, Bethesda, MD). Secondary LICOR goat anti-rabbit IRDye® 800 (Licor, Cat #926-32211) or goat anti-mouse IRDye®680 (Licor, Cat #926-68070) antibodies were diluted at 1:5,000 in TBS-T and incubated at room temperature for 1 hour. All membranes were imaged on LICOR Odyssey infrared imaging system at 680 and 800nm wavelengths and analyzed with ImageStudio Odyssey Lite Software (LICOR).

#### Incucyte Caspase 3/7 assay

LN443-Cas9 cells were transduced with guide RNA sgCtrl or sgVRK1. Following 1 wk under selection, 5×10^4^ cells per well were seeded in 24-well Falcon plates (Fisher Scientific, Cat #353047). 5 mM of IncuCyte Caspase-3/7 Green Apoptosis Assay Reagent (Sartorius, cat #4440) as well as 1:500 of Nuclight Rapid Red Dye (Sartorius, cat #4717) were added to each well. The plate was transferred into the IncuCyte S3 Live-Cell Analysis System (Sartorius, cat #4647) for imaging. Phase contrast images and green/red fluorescent channel images were captured using the 10x objective magnification every four hours for a total of 48 hours. For each well, four images containing both phase contrast and green channel data were obtained.

Using the IncuCyte S3 Analysis System software, cell confluence over time was quantified along with the intensity of green (apoptosis positive) objects in mm2/well. Computer generated masks for confluence and green area, trained on a sample set of images across time points and confluency levels, were manually checked for accuracy. Each metric was averaged over the four quadrants per well. First, the green object total intensity metric for each well was divided by the confluence metric for each well, yielding a normalized measure of Caspase-3/7 activity.

#### Mass Spectrometry Sample Preparation

Samples were processed with the SL-TMT protocol and phospho-enrichment methods described previously (42). Data were acquired with Orbitrap Eclipse mass spectrometer with FAIMS and coupled to a Proxeon NanoLC-1200 UHPLC (ThermoFisher Scientific).

#### Mass Spectrometry Data Analysis

A suite of in-house software tools were used for .RAW file processing and controlling peptide and protein level false discovery rates, assembling proteins from peptides, and protein quantification from peptides as previously described (43). MS/MS spectra were searched against a Uniprot Human database with both the forward and reverse sequences. Database search criteria are as follows: tryptic with two missed cleavages, a precursor mass tolerance of 50 ppm, fragment bin tolerance of 0.02, static alkylation of cysteine (57.02146 Da), static TMT labeling of lysine residues and N-termini of peptides (304.2071 Da), variable oxidation of methionine (15.99491 Da) and variable phosphorylation on serine, threonine, and tyrosine (+79.966 Da). Phosphorylation site localization was determined using the AScore algorithm (44) using a threshold of 13 corresponding to 95% confidence in site localization. TMT reporter ion intensities were measured using a 0.003 Da window around the theoretical m/z for each reporter ion. Proteins with <100 summed signal-to-noise across all channels and <0.5 precursor isolation specificity were excluded from the final dataset.

Ratios were calculated between peptide quantitation at 4hrs post-dTAG^V^-1 versus 0hrs and at 8hrs post-dTAG^V^-1 versus 0hrs. P-values for each ratio were calculated using Student’s T-test. From the fold change values and p-values, Kinase-Substrate Enrichment Analysis (KSEA; https://casecpb.shinyapps.io/ksea/) was performed with NetworKIN score cutoff of 3 (14,45–47).

#### Immunofluorescence

The nuclear membrane and DNA damage foci were visualized by immunofluorescence using the following procedure. LN443-Cas9, SF172-Cas9, NB-1, and Kelly cells were transduced with various sgRNAs. BAF and Emerin were imaged by transducing GFP-tagged constructs. EGFP-BAF was a gift from Daniel Gerlich (Addgene plasmid #101772) (48). pLVX-EF1a-EGFP-Emerin-IRES-Hygromycin was a gift from David Andrews (Addgene plasmid #134864) (49). For the BAF experiment, inducible BAF wild-type or mutant expression vectors were transduced in LN443 cells. Following selection (~5-7days), doxycycline induction (0.5μM for 3 days), or dTAG^V^-1 treatment (0.5μM for 1 day), cells were seeded onto #1½ cover-glasses (Sigma, Cat #CLS285018) in 6-well Falcon plates (Fisher Scientific, Cat #087721B). The next day, cells were fixed with 4% Formaldehyde (VWR, Cat# 100503) diluted in PBS. Fixed cells were permeabilized and blocked with 0.1% Triton-X in 50% Odyssey Blocking Buffer (Licor, Cat# 927-70001) in PBS for 1hr at room temperature. The cells were then incubated with the primary antibody at the specified dilution in 0.1% Triton-X with 50% Odyssey Blocking, overnight at 4°C. After washing three times with PBS, cells were incubated with the secondary antibody at the specified dilution in 0.1% Triton-X with 50% Odyssey Blocking for one hour at room temperature. The cells were then washed three times with PBS and mounted onto glass slides with ProLong Gold anti-fade mounting media with DAPI (Life Technologies, Cat #P36941). Imaging was conducted using an Olympus IX73 inverted microscope, an Olympus DP80 CCD camera, and 20x/40x/100x objectives. Antibodies used were the following:

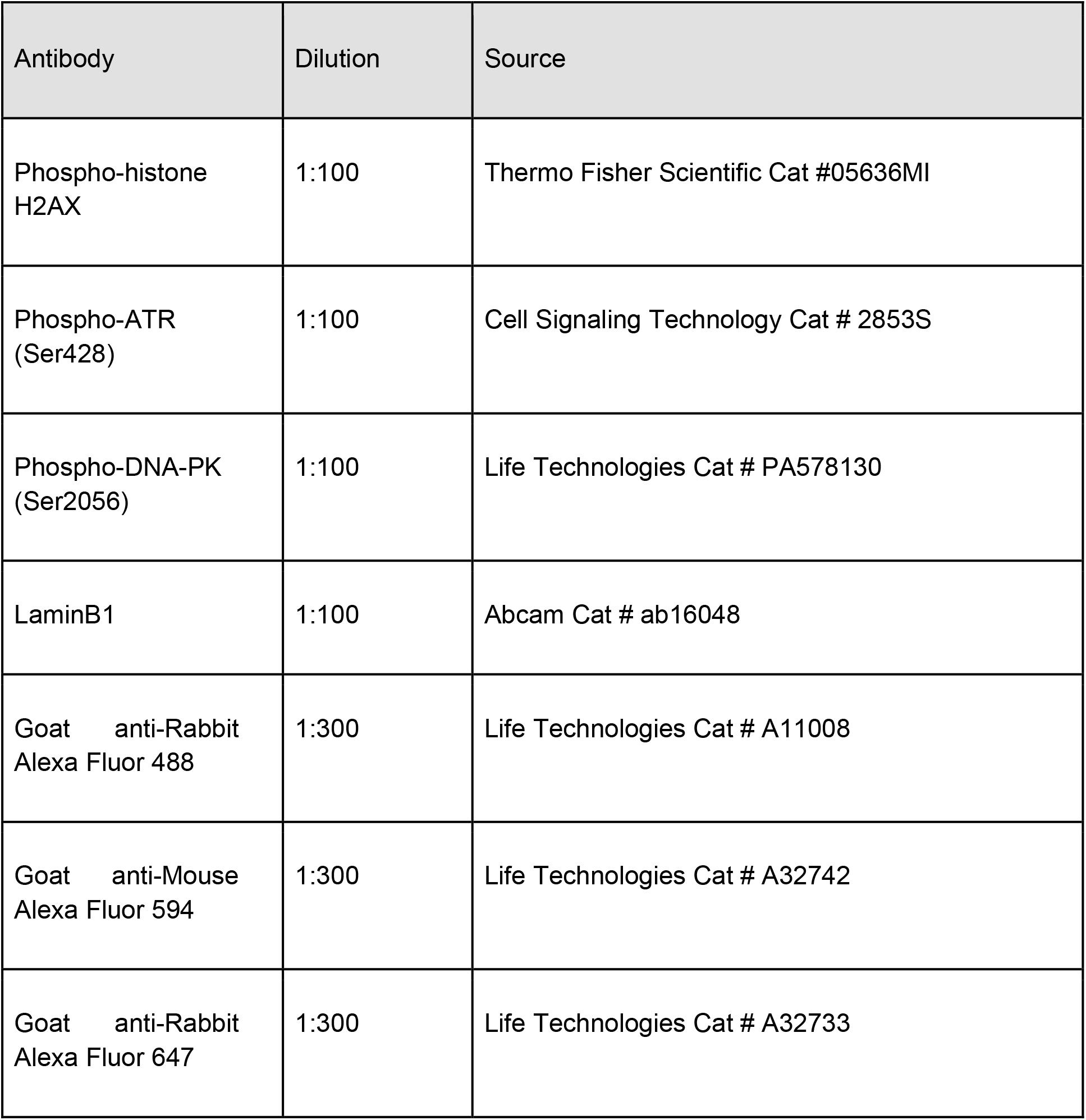

| Antibody | Dilution | Source |
| --- | --- | --- |
| Phospho-histone H2AX | 1:100 | Thermo Fisher Scientific Cat #05636MI |
| Phospho-ATR (Ser428) | 1:100 | Cell Signaling Technology Cat # 2853S |
| Phospho-DNA-PK (Ser2056) | 1:100 | Life Technologies Cat # PA578130 |
| LaminB1 | 1:100 | Abcam Cat # ab16048 |
| Goat anti-Rabbit Alexa Fluor 488 | 1:300 | Life Technologies Cat # A11008 |
| Goat anti-Mouse Alexa Fluor 594 | 1:300 | Life Technologies Cat # A32742 |
| Goat anti-Rabbit Alexa Fluor 647 | 1:300 | Life Technologies Cat # A32733 |

#### Live-cell Imaging

dTAG-VRK1-LN443 or dTAG-VRK1-NB-1 cells were transduced with EGFP-BAF as previously described. 2×10^4^ - 5×10^4^ per well were then seeded in MatTek 24-well, glass-bottom plates (Fisher Scientific NC1284979). 4 hours following dTAG^V^-1 addition, the plate was imaged using the 40x objective in a Leica DMi8 Widefield microscope with automated stage, an Oko-Lab stage-top incubator, and Oko-Lab CO2/Humidity controller. 3×3 fields per well were imaged every 20min for 48hrs. Image stitching was performed using the Leica LAS X software platform. Subsequent image analysis was performed using ImageJ ver. 1.53m.

#### *In vivo* inducible sgRNA xenografts

This study was approved by the Institutional Care and Use Committee (IACUC) of Dana-Farber Cancer Institute and performed under protocol 15-029. IACUC guidelines on the ethical use and care of animals were followed. SF295 cells constitutively expressing Cas9 were infected with tamoxifen-inducible sgRNAs targeting Chr2-2 or *VRK1*. 6.0e6 cells were resuspended in 2:1 vol/vol matrigel:media and subcutaneously implanted into the left and right fat pads of 6-to-8-week old female NSG (NOD-*scidl* IL2Rg^null^) mice (Jackson Laboratory stock no. 005557). When either tumor was ~100-200 mm^3^ mice were randomized to tamoxifen or vehicle treatment. Tamoxifen was delivered by 3 daily intraperitoneal injections of ~3mg. Tamoxifen (Sigma-Aldrich) was prepared at a stock concentration of 30mg/mL in corn oil. The control group received an equal volume of corn oil. Tumors were measured by Vernier caliper and volume was determined using the standard formula [(length x width^2^)/2 where length is always the larger measurement]. Animals were euthanized once they reached a humane endpoint and tumor tissue was flash frozen or formalin fixed for later protein and genomic DNA extraction. All mice that developed tumors were included in the analysis.

7 days following treatment with tamoxifen or vehicle control, tumors were collected from a subset of xenografted mice. These were fixed in formalin and embedded in paraffin. Immunohistochemistry was performed following standard protocol, staining for phospho-H2AX (S139) (CST cat# 9718; 1:500).

#### Data Availability Statement

The mass spectrometry proteomics data have been deposited to the ProteomeXchange Consortium via the PRIDE (50) partner repository with the dataset identifier PXD030599.

