## Supplemental figures 1-8 for "VRK1 is required in VRK2-methylated cancers of the nervous system"

### Supplementary Figure Legends

**Supplementary figure 1. Validation of *VRK1* CRISPR KO sgRNAs in GBM, NB, and DMG models**

**A.** Immunoblot of VRK1 protein expression following expression of 2 different sgRNAs in the DMG cell lines BT869Luci and SU-DIPGXIIIP*Luci. sgChr2.2 and sgLacZ served as cutting and non-cutting controls, respectively.

**B.** Immunoblot showing VRK1 expression in LAN-1, SK-N-BE(2)C, or Kelly cell lines with sgRNAs targeting either sgCtrl or *VRK1*.

**C.** Immunoblot showing VRK1 expression in SF172 or LN443 GBM cell lines with sgRNAs targeting either sgCtrl or *VRK1*.

**Supplementary Figure 2. Cell cycle distribution of GBM, NB, and DMG models following VRK1 depletion**

**A.** Cell cycle distribution of Kelly cells 7 days following *VRK1* KO in three independent sgRNAs. Significance was determined by one-way ANOVA within each phase of cell cycle. * $p < 0.05$, ** $p < 0.001$, *** $p < 0.0001$

**B.** Cell cycle distribution of BT869Luci DMG cells 7 days following *VRK1* KO. Significance was determined by Student's T-test. * $p < 0.05$

**C.** Representative cell cycle distribution from propidium iodide staining of SF172 GBM cells following combinations of sgCtrl/Ctrl, sgCtrl/VRK1, sgVRK2/Ctrl, or sgVRK2/VRK1 guides at 7 days post-infection. Cell cycle analysis was performed using FlowJo (ver.10.8.0).

**D.** Representative cell cycle distribution from propidium iodide staining of LN443 GBM cells 7 days following *VRK1* KO. Cell cycle analysis was performed using FlowJo (ver.10.8.0).

**Supplementary Figure 3 VRK1 and VRK2 expression in normal tissues**

**A.** Violin plots showing Log2(TPM+1) mRNA expression for *VRK1* (top panel) and *VRK2* (bottom panel) across a panel of healthy tissues from GTEx (https://gtexportal.org/).

#### Supplementary Figure Legends continued

**Supplementary Figure 4 VRK1 and VRK2 expression in GBM and NB tumors**

**A-B.** UMAP plots from Cellligner-corrected *VRK2* (A) or *VRK1* (B) expression for RNA-sequencing on all available human tumors. Brain tumor (red box) and neuroblastoma (blue box) tumor lineage clusters are indicated with boxes. Right panels: violin plots showing Cellligner-corrected *VRK2* or *VRK1* expression for all tumor lineages (black) against brain cancers and NB (red).

**C.** Violet plots showing log2(TPM) RNA-sequencing data from the TREEHOUSE/TARGET dataset containing human neuroblastoma tumors. Tumors were separated on the basis of MYCN-amplification (top panel) or adrenergic/mesenchymal (bottom panel).

**D.** Immunoblot of basal protein levels of VRK1 and *VRK2* in a panel of pediatric H3.3K27M and H3 wild-type glioma cell lines.

*p < 0.05, **p < 0.001, ***p < 0.0001; Two-tailed, Student's T-test for all comparisons.

**Supplementary Figure 5. Validation of paralog relationship of VRK1 and VRK2 through VRK2 depletion or over-expression**

**A.** Immunoblot of basal protein levels of VRK1 and VRK2 in a panel of GBM cell lines.

**B.** Immunoblot of VRK2 protein expression following generation of isogenic cell line pairs in *VRK2*$^{high}$ GBM cell lines through CRISPR KO of *VRK2*.

**C.** Incucyte time-lapse experiment of cell proliferation in SF172 cell line expressing 2x2 combinations of sgCtrl/Ctrl, sgCtrl/VRK1, sgVRK2/Ctrl, or sgVRK2/VRK1 guides. t = 0hrs is 7 days post-infection under antibiotic selection. Significance at each time point was determined by two-way ANOVA (treatment x time). * p <0.05.

**D.** Immunoblot of exogenous wildtype or kinase-inactive VRK1$^{WT}$, VRK1$^{K179E}$, VRK2$^{WT}$, or VRK2$^{K168E}$ following lentiviral transduction of LN443 and LNZ308 GBM cell lines.

**E.** Clonogenic assay of LN443 or LNZ308 GBM cell lines overexpressing VRK2$^{WT}$ or kinase-inactive VRK2$^{K168E}$ 3 weeks following lentiviral transduction with non-targeting guide or guides targeting *VRK1* (sgVRK1#1 and sgVRK1#2).

**F.** Clonogenic assay of the NB-1 neuroblastoma cell line overexpressing VRK2$^{WT}$ or kinase-inactive VRK2$^{K168E}$ 2 weeks following lentiviral transduction with sgCh2.2 control guide or sgVRK1 guide.

**G.** Immunoblot of protein expression levels of exogenous VRK1$^{WT}$, VRK1$^{K179E}$, VRK2$^{WT}$, or VRK2$^{K168E}$ following lentiviral transduction of SU-DIPGXIIIP*Luci cells.

**H**. Effect of VRK2$^{WT}$ or VRK2$^{K168E}$ overexpression on SU-DIPGXIIIP*Luci cell viability following 10 days *VRK1* KO. (n=4; mean ± SD).

*p < 0.05, **p < 0.001, ***p < 0.0001; Two-tailed, Student's T-test

#### Supplementary Figure Legends continued

**Supplementary Figure 6. dTAG degrader system for ligand-induced VRK1 depletion**

**A.** Left panel: Immunoblot validation of the dTAG-VRK1-dTAG degrader system in Kelly neuroblastoma cells. Exogenous dTAG-VRK1-dTAG was degraded in the presence of $dTAG^{V}$-1 (0.75 µM). Endogenous *VRK1* was independently targeted with CRISPR KO. sgNT is a non-targeting guide control. Right panel: Viability of dTAG-VRK1-Kelly cells following addition of either vehicle control or 0.75 µM $dTAG^{V}$-1. Significance at each time point was determined by two-way ANOVA (treatment x time). * p <0.05

**B.** Incucyte time-lapse experiment of cell proliferation in dTAG-VRK1-LN443 cells following transduction of non-targeting guide (sgCtrl) or guide targeting *VRK1* (sgVRK1). t = 0hrs is 5 days post $dTAG^{V}$-1 (0.5 µM) addition. Significance at each time point was determined by two-way ANOVA (treatment x time). ** p <0.001.

**Supplementary Figure 7. Phospho-peptide quantification of LEM-domain containing proteins following acute VRK1 depletion**

**A.** Change in phospho-peptide abundance at 4h and 8h following VRK1 degradation in dTAG-VRK1-NB-1 cells. Each point represents a separate phospho-peptide measured by quantitative phospho-proteomics. Highlighted are three nuclear membrane associated, LEM-domain containing proteins (TMPO, LEMD3, and EMD).

**Supplementary Figure 8. Nuclear morphology changes following VRK1 and VRK2 depletion via decreased BAF phosphorylation**

**A.** Nuclear membrane morphology in the SF172 GBM cell line following transduction with 2x2 combinations of sgCtrl/Ctrl, sgCtrl/VRK1, sgVRK2/Ctrl, sgVRK2/VRK1 guides. Nuclear membrane was visualized by immuno-fluorescent staining for LaminB1. White arrow points to nuclear bridge. Blue arrow points to micro-nuclei.

**B.** Immunoblot of phosphorylated BAF (S4) and total BAF following 5 days doxycycline-induced expression of guide targeting *VRK1* in SU-DIPGXIIILuci DMG neurospheres. Representative of 2 independent experiments. pBAF and total BAF were probed in two separate blots of the same lysate.

**C.** Immunoblot following 3 days of doxycycline-induced expression of $BAF^{WT}$, $BAF^{S4A}$, $BAF^{S4D}$ in LN443 GBM cells.

**D.** Time-lapse of live-cell experiment showing nuclear envelope morphology (GFP-tagged BAF) following VRK1 degradation in dTAG-VRK1-NB-1 NB cells undergoing mitosis ($dTAG^{V}$-1 0.5µM).

### Supplementary Figure 1.

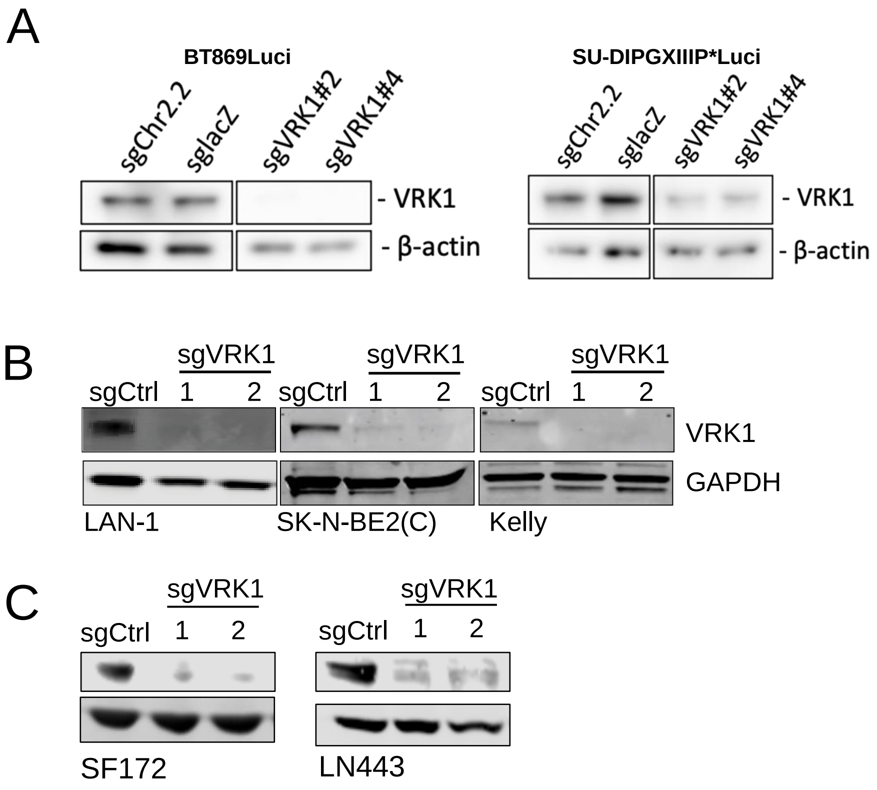

Supplementary Figure 2.

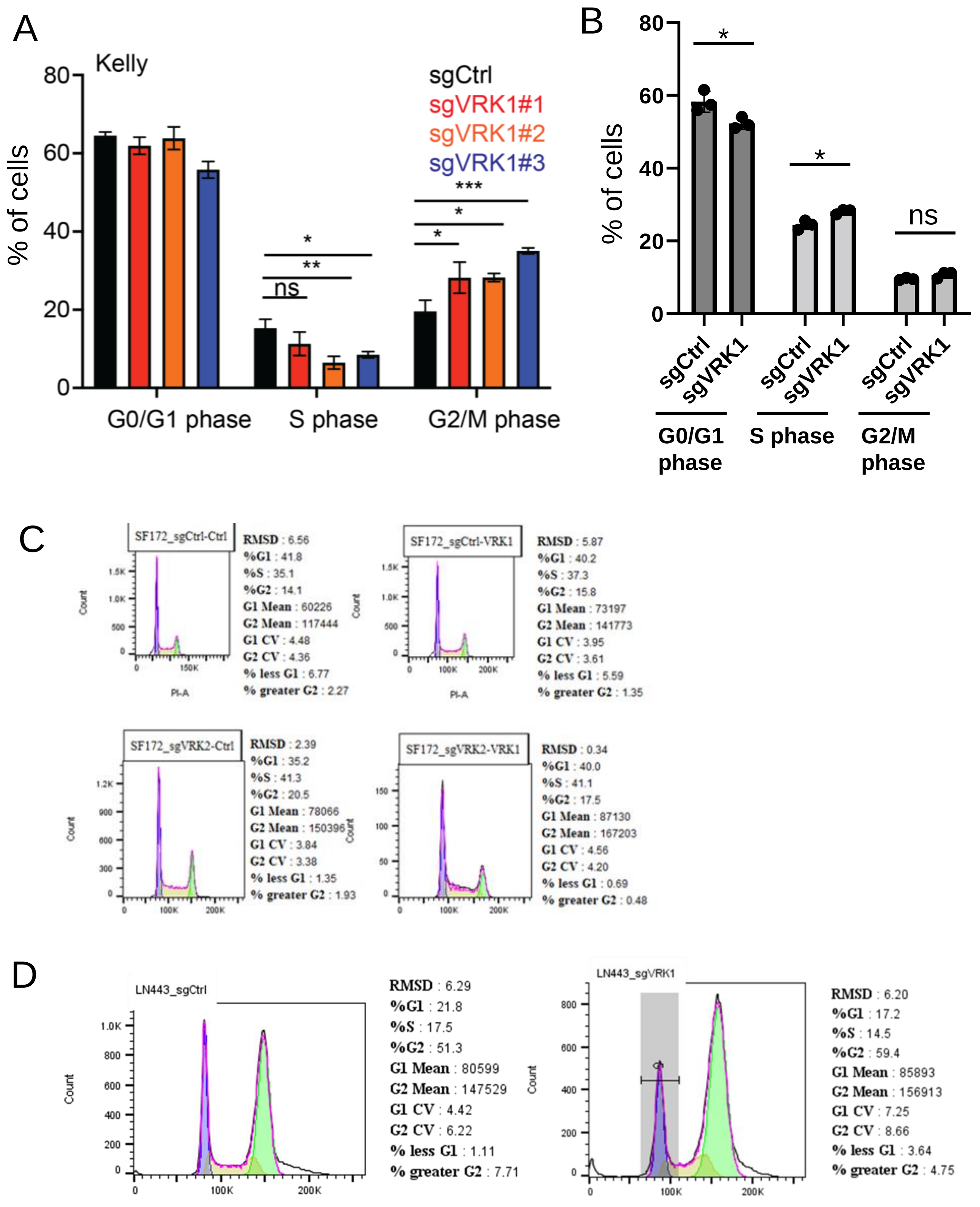

Supplementary Figure 3.

A

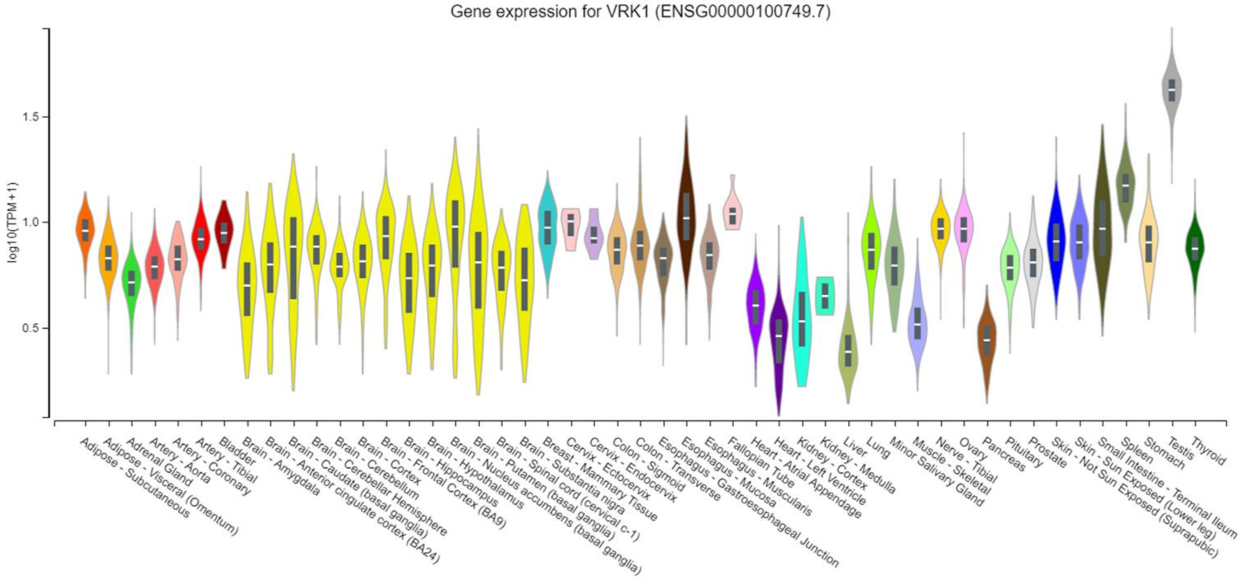


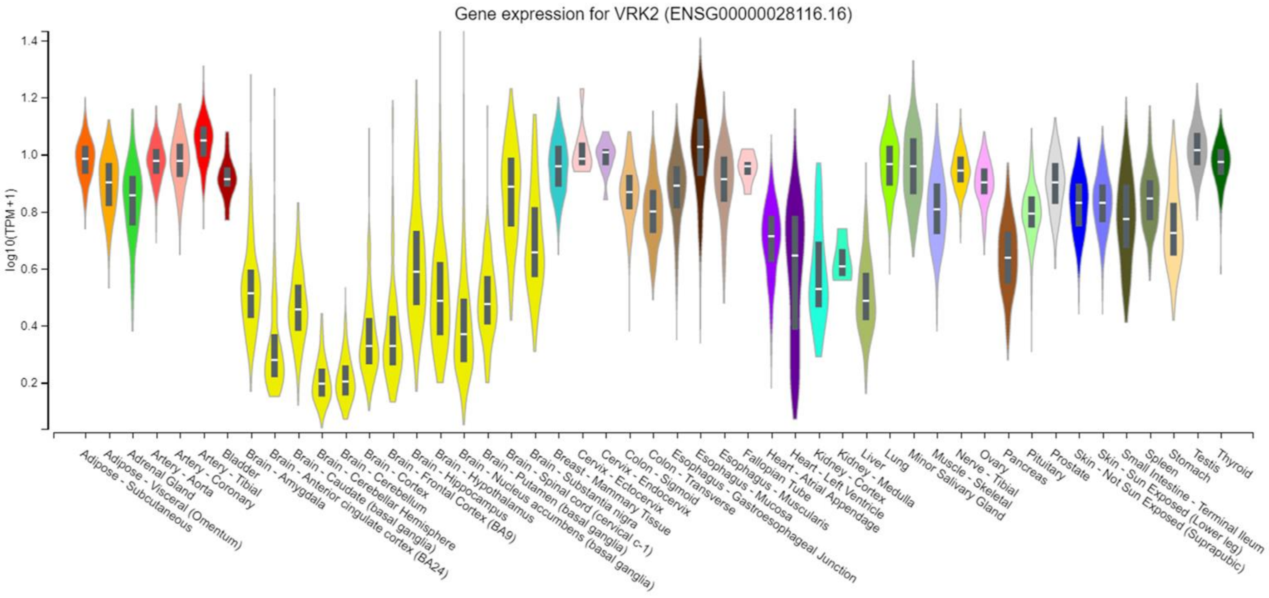

### Supplementary Figure 4.

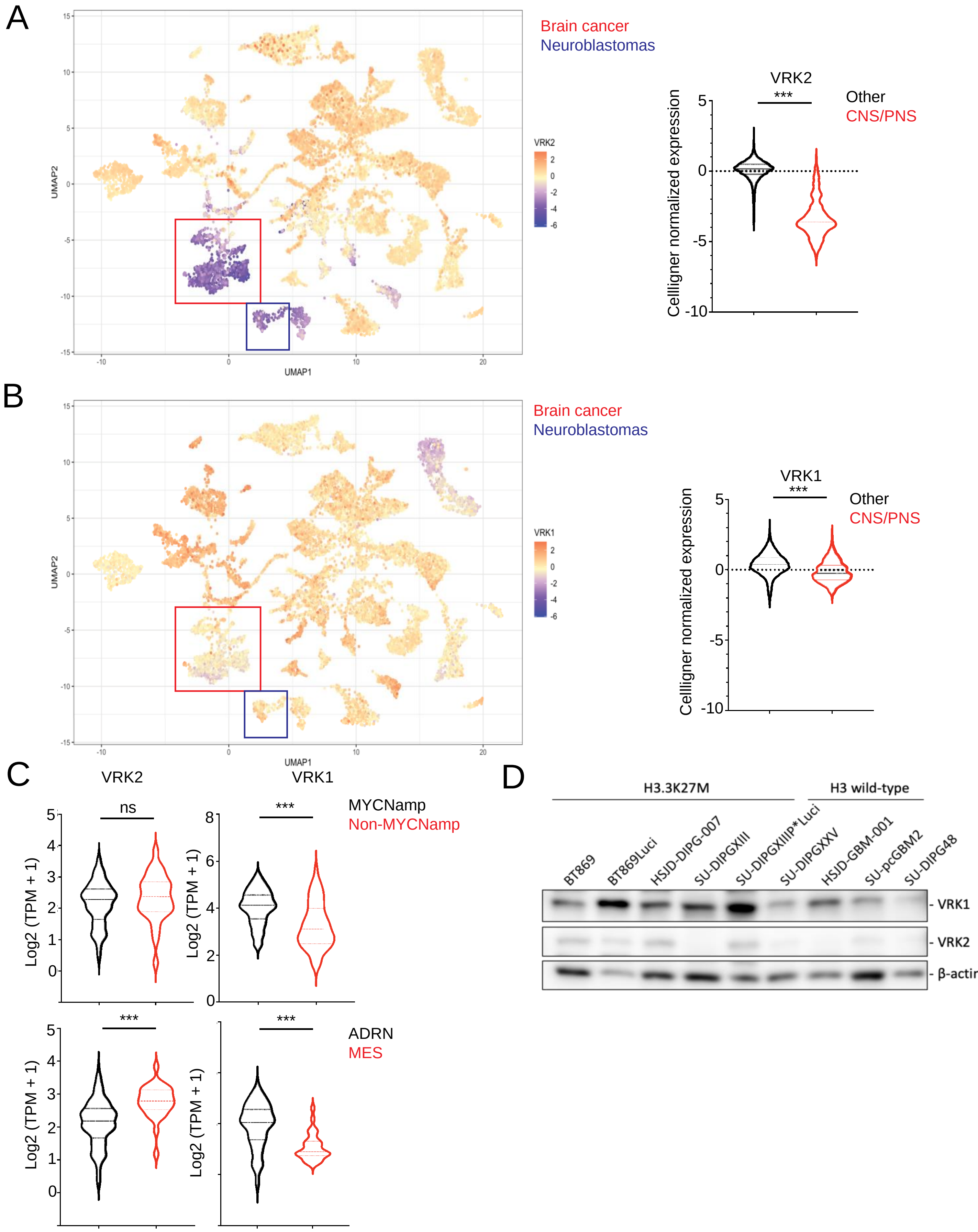

Supplementary Figure 5.

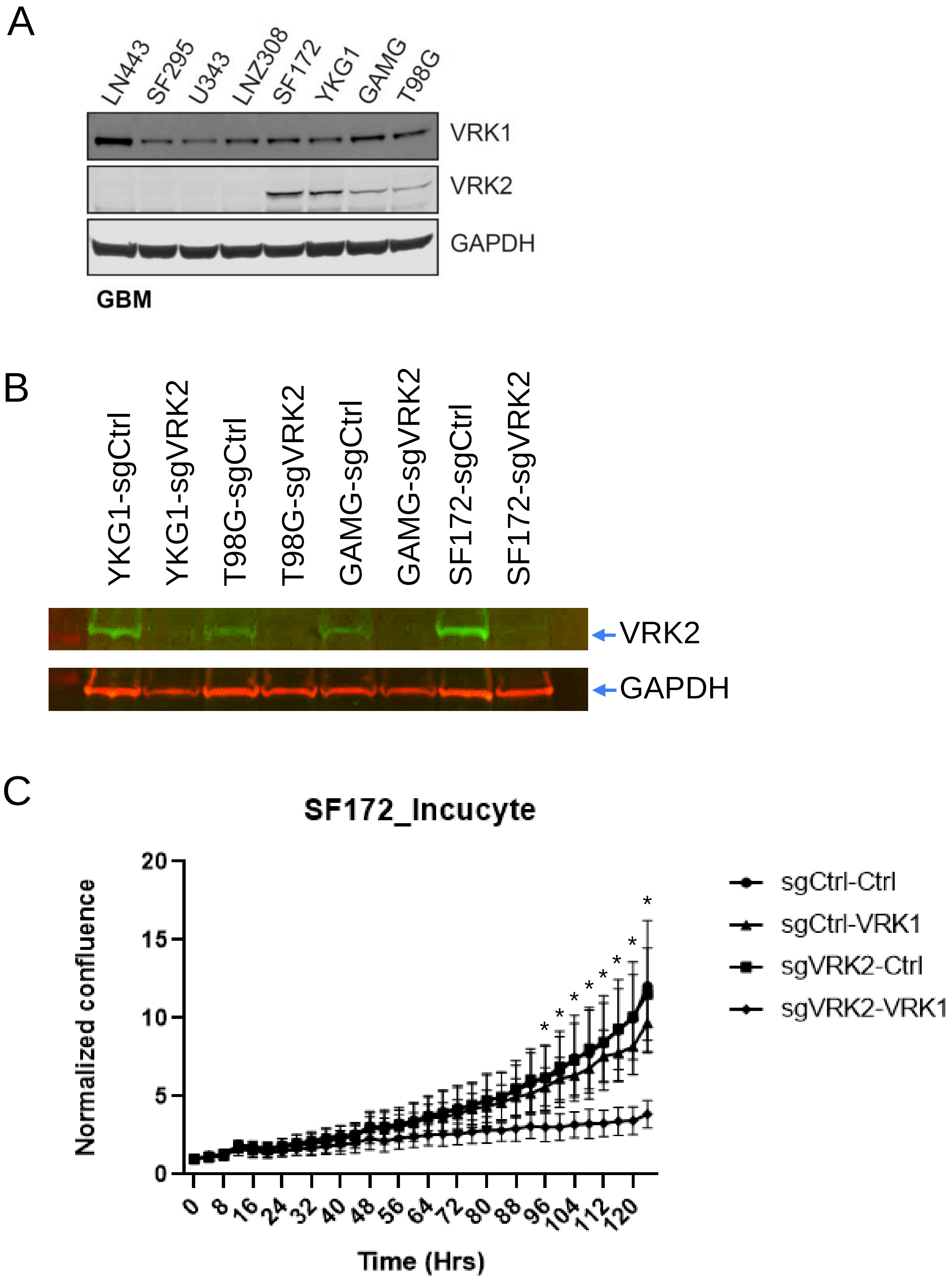

### Supplementary Figure 5 cont.

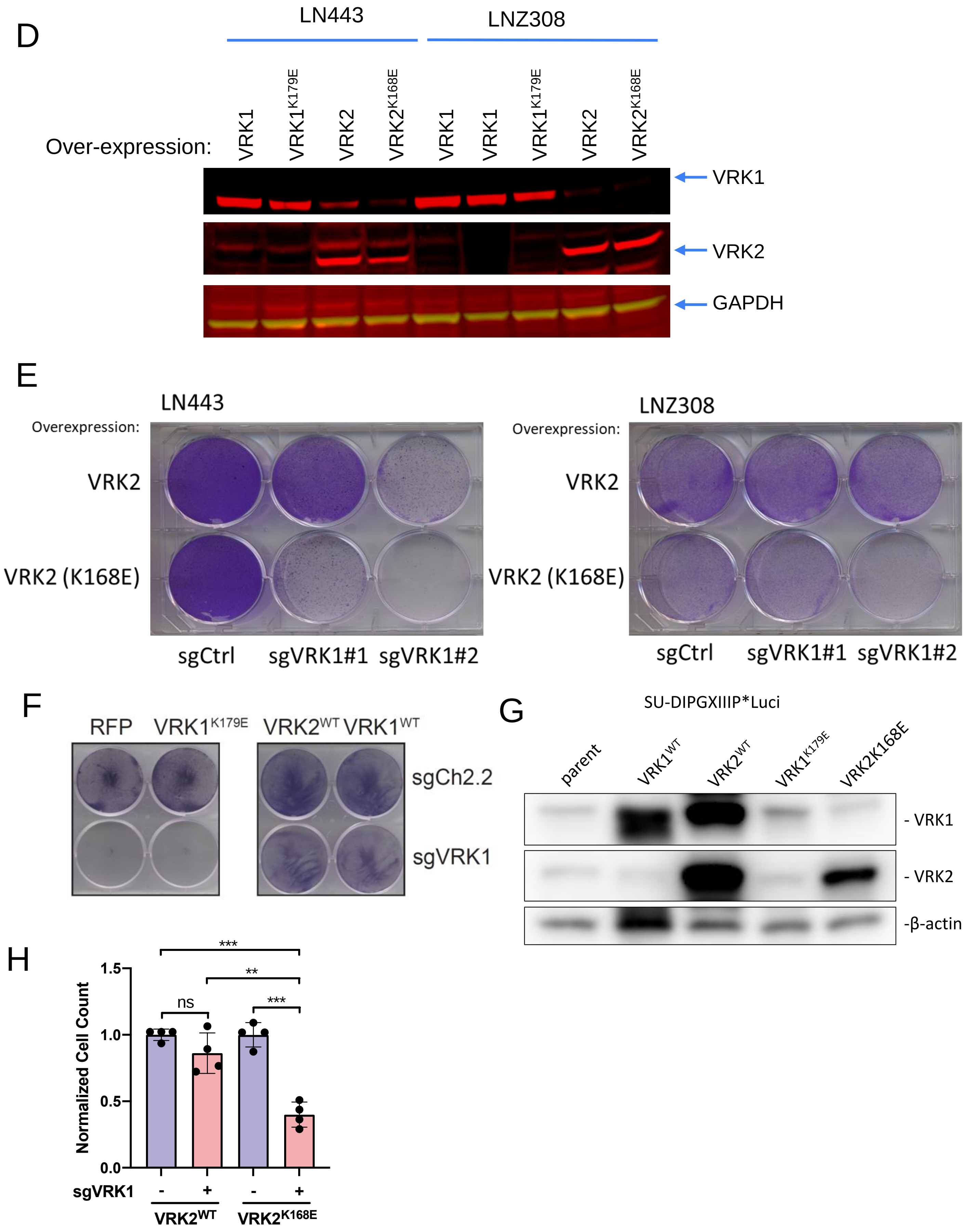

Supplementary Figure 6.

A

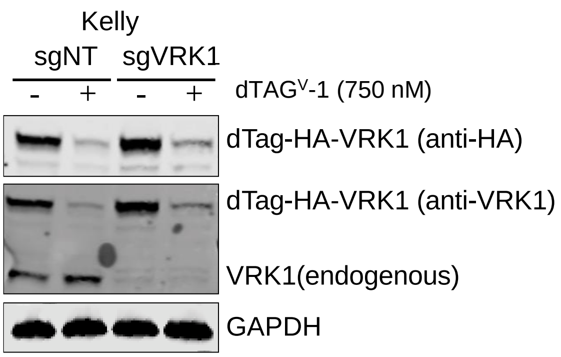


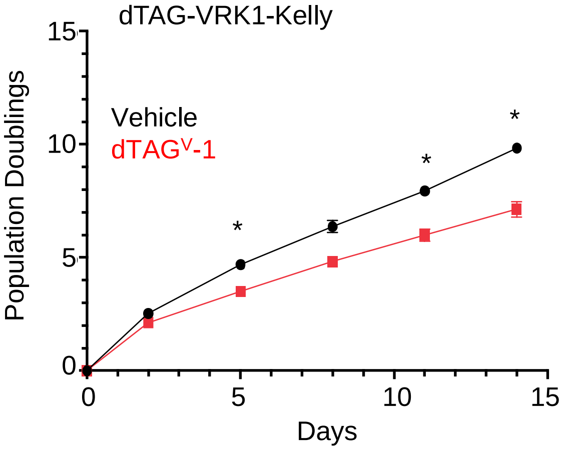


B

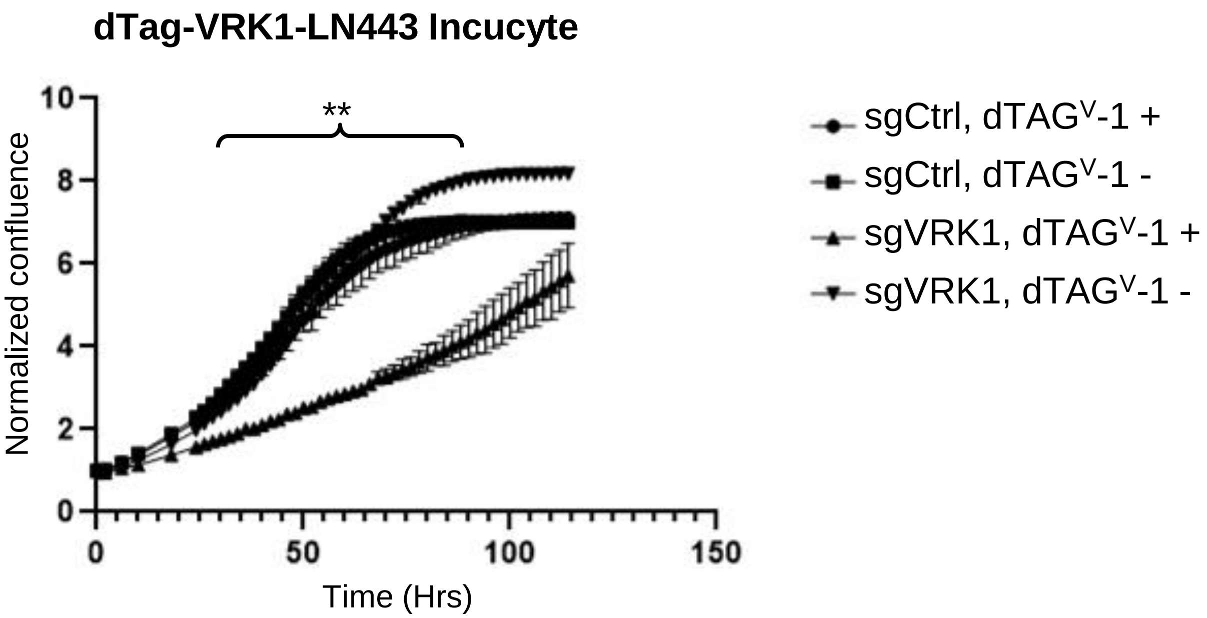

Supplementary Figure 7.

A

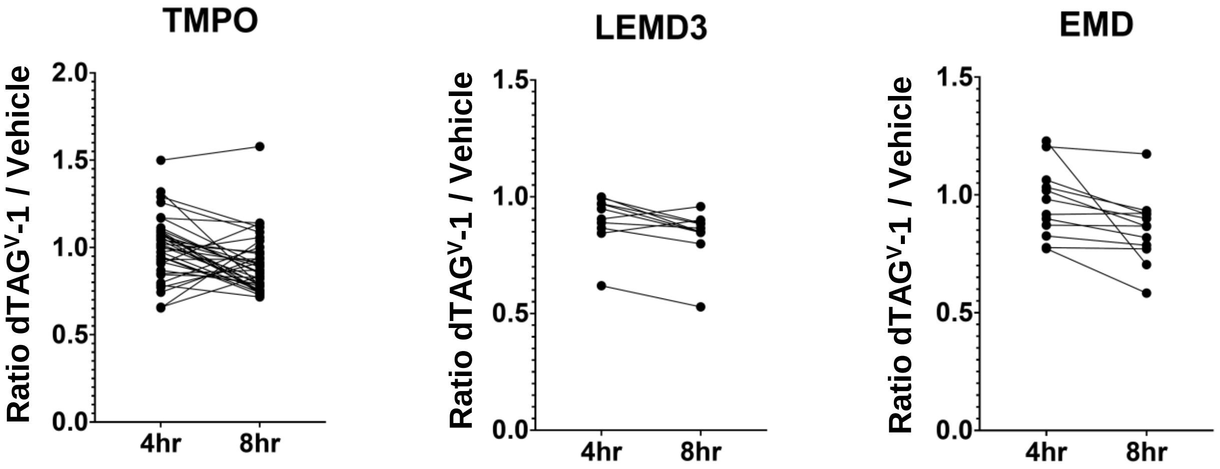

### Supplementary Figure 8.

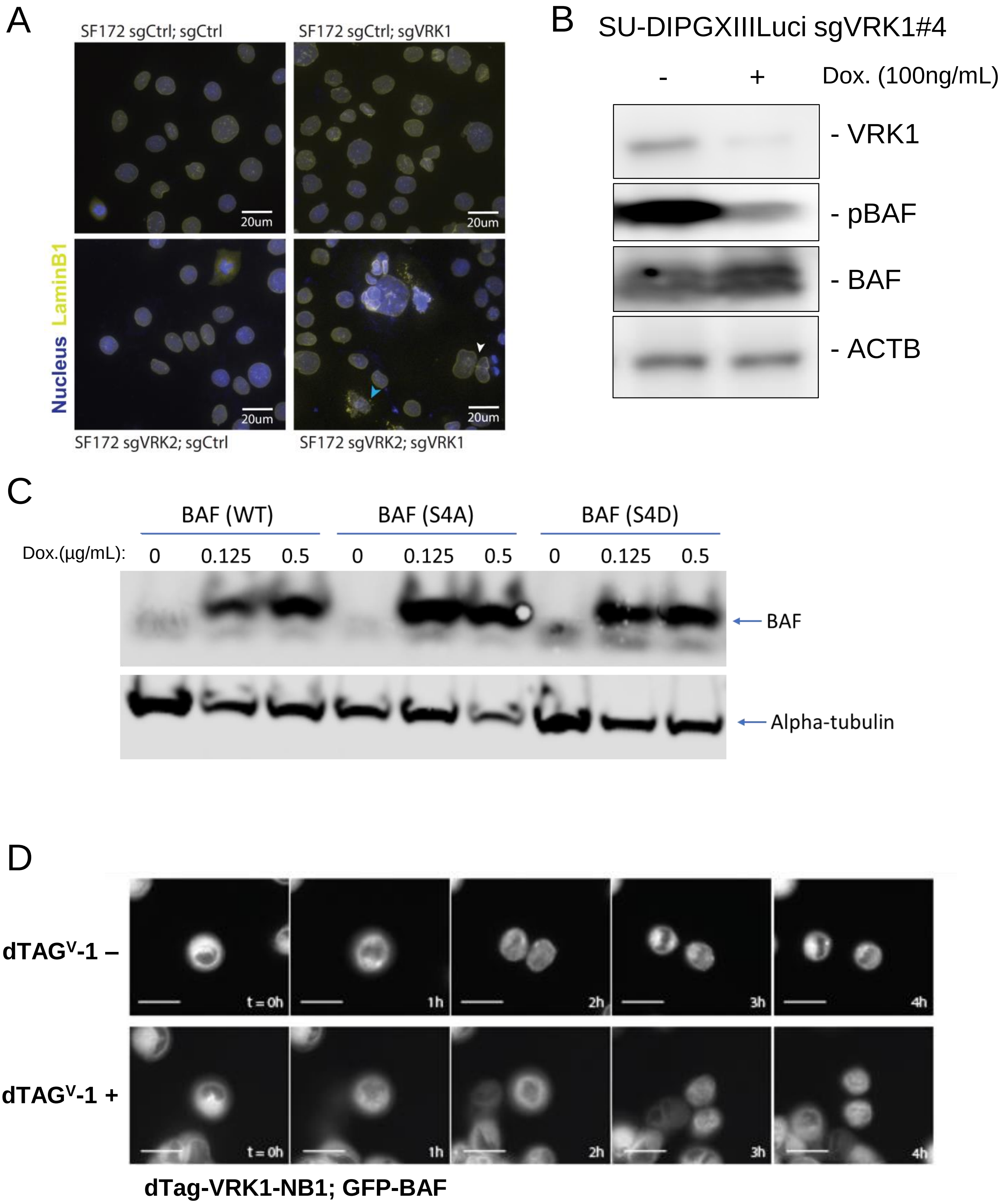
